# Constant light impairs memory processing transgenerationally in *D. melanogaster*

**DOI:** 10.1101/2025.06.04.657854

**Authors:** Rebecca S. Moore, Amita Sehgal

## Abstract

Environmental perturbations can have profound effects on our physiology and behavior, but their long-lasting impact remains debatable. We discovered that changes in standard light-dark conditions, such as exposure to constant light or simulated chronic jetlag, causes loss of associative memory in appetitive and aversive conditioning paradigms in *D. melanogaster*, and this behavior persists through three generations despite transfer of progeny to a standard light-dark cycle. Impaired memory is transmitted through females and is independent of any changes in fitness, brain or mushroom body architecture, or sensing acuity. Mechanistically, effects of constant light on memory are mediated by increased PIWI/piRNA pathway expression in the germline and its H3K9me3 writing capabilities, acting through altered expression of the Dopamine-1-like Receptor 1 (Dop1R1) in the brain. These findings suggest that irregular exposure to light, such as nighttime light pollution, can have negative consequences across generations.

**HIGHLIGHTS:**

- Constant light impairs associative memory in Drosophila for three generations
- *piwi* is required in the brain and germline for inheritance
- Dopamine 1 Receptor 1 activity is dampened by constant light

## INTRODUCTION

Light is an essential environmental factor to almost all life forms. In particular, the natural light-dark cycle plays a critical role in synchronizing behavioral and biological processes, typically through the circadian clock. However, mistimed or excessive exposure to light can negatively affect physiology and behavior, for instance cognition. In humans, cognitive function varies over the 24 h day; while the exact timing can differ from person to person, it typically starts off low in the morning and increases throughout the day, staying at high levels until habitual bedtime, apart from a dip in the afternoon (Adan et al., 2012; Horne and Ostberg, 1976; Schmidt et al., 2007). Large scale epidemiological studies have shown that deviations in the natural light-dark cycle affect associative learning and memory, and imaging studies have demonstrated impacts of light exposure on cortical and subcortical networks involved in cognitive processes, such as attention, arousal, and memory (Chellappa et al., 2011; Perrinet, 2004; Vandewalle et al., 2011, 2007).

At the same time, small model organisms have been used to study the molecular and cellular effects of mistimed light exposure. For example, adult mice and rats or hamsters subjected to chronic light or repeated phase shifts in light exposure have impairments in learning and memory ability or reduced neurogenesis respectively (Fujioka et al., 2011; Ma et al., 2007; Gibson et al., 2010;). In *Drosophila,* long term memory is impaired when flies are reared in constant darkness due to inactivity of the transcription factor cAMP response element-binding protein (CREB) (Inami et al., 2020). Together, these studies show that regular light-dark cycling is essential to proper behavioral performance. However, modern lifestyles usually do not incorporate appropriate exposure to light:dark cycles. Indoor lighting does not substitute for sunlight during the day, and, importantly, artificial light at night results in an almost constant light environment.

Over the past decade, emerging evidence indicates that effects of the environment can also be transmitted to subsequent generations, a phenomenon known as transgenerational epigenetic inheritance (TEI). For example, male mice trained to associate fear with an odor transmit odor-sensitivity to their sons (Dias and Ressler, 2014). Researchers concluded that offspring possessed an increased abundance of sensory neurons specific to the same odor their fathers were trained to fear (Dias and Ressler, 2014). On the other hand, environmental enrichment activities can ameliorate behavioral defects of mice defective in long-term potentiation and memory, and the rescue is heritable through the activation of p38 signaling (Arai et al., 2009). In many cases, mechanisms underlying TEI are not known, but where examined, they mostly implicate DNA methylation, histone modification, and non-coding RNAs (Perez and Lehner, 2019). In *C. elegans* wild-type worms exposed to pathogenic *P. aeruginosa* learn to avoid it and switch their feeding behavior to nonpathogenic bacteria. This behavior is transgenerationally inherited to naïve progeny up to the fourth generation through transforming growth factor β (TGF-β) signaling in sensory neurons and the PIWI Argonaute small RNA pathway (Moore et al., 2019). A different mechanism explains why exposure of *D. melanogaster* to predatory wasps leads up to five future generations of flies to preferentially lay their eggs in ethanol-rich food to prevent wasp infection; this effect is mediated by Neuropeptide-F (NPF) inhibition in the brain leading to germline reprogramming through caspase activity (Bozler et al., 2019). These few examples suggest that parental environment can have profound impacts on subsequent generations. Elucidating mechanisms underlying these environmentally triggered epigenetic programs is essential for understanding the fundamental principles upon which biological inheritance is based.

*Drosophila melanogaster* has emerged as a premier organism to study TEI, due to its short generation time, large broods, and evolutionarily conserved epigenetic mechanisms that encode and transmit transgenerational information. In this study, we found that deviations in the normal light-dark cycle (i.e. constant light or chronic jetlag) impair the standard behavioral response in appetitive and aversive associative short- and long-term memory paradigms. Not only is memory performance impacted in the parental generation, but progeny raised in a normal light-dark cycle inherit this impaired memory for three generations before returning to baseline in the fourth. The mechanism involves the PIWI-interacting RNA pathway, which acts via H3K9me3 epigenetic marks, in the germline and brain and alterations in Dopamine 1 Receptor 1 activity in the brain. Inheritance is through the female germline and independent of changes in brain architecture, reward sensing, or fitness. These results suggest that the negative behavioral consequences of exposure to irregular light cycles persist beyond the parental generation to have long lasting effects on future generations.

## RESULTS

### Raising flies in constant light does not affect sleep or behavioral rhythms in adults

As noted above, constant light can be deleterious to health, so we asked if it could have long lasting effects. Using a Drosophila model, we raised flies in constant light (LL) for three generations and then tested adults for sleep and behavioral rhythms. LL-rearing did not affect sleep in males or females maintained subsequently in a light:dark cycle (Figure S1A). Nor did it affect daytime (ZT0-12) or nighttime (ZT12-24) waking activity levels, which were comparable to those of flies raised in LD (Figure S1B). Likewise, free-running circadian rhythms were unimpaired. Male or female flies raised in constant light and tested for locomotor activity rhythms in total darkness did not show any significant difference from light:dark reared flies (Figure S1C).

### Constant light or a jetlag paradigm abrogates associative memory

Although LL did not have lasting effects on sleep and circadian rhythms, it could impact other behaviors. Drosophila maintained in a standard 12:12 light dark (LD) cycle can be trained to associate an odor with a sugar reward (CS+), or to avoid an odor when paired with an electric shock (CS+) (Quinn et al., 1974; Tully and Quinn, 1985). To test whether changes to the standard LD cycle would affect short- or long-term appetitive memory, we raised wild-type (Canton S, red eyed) flies in either standard LD or constant light (LL, 24 hour/day) for at least 3 generations and then assayed for both appetitive and aversive associative memory in constant darkness with a red light (Figure 1A). We found that chronic exposure to constant light abrogates short-term appetitive memory (Figure 1B). Flies were re-fed for three hours on the bench and then starved again overnight in LD or LL and then tested for long-term appetitive memory. Like short-term memory, long-term memory was impaired in flies raised in LL compared to LD (Figure 1C). Short- and long-term aversive memory was similarly affected by LL (Figure 1D-E). In subsequent experiments, we discovered that one generation of constant light exposure was sufficient to impair appetitive memory formation, like 3 generations of LL (Figure S1D-E).

**Figure 1:**
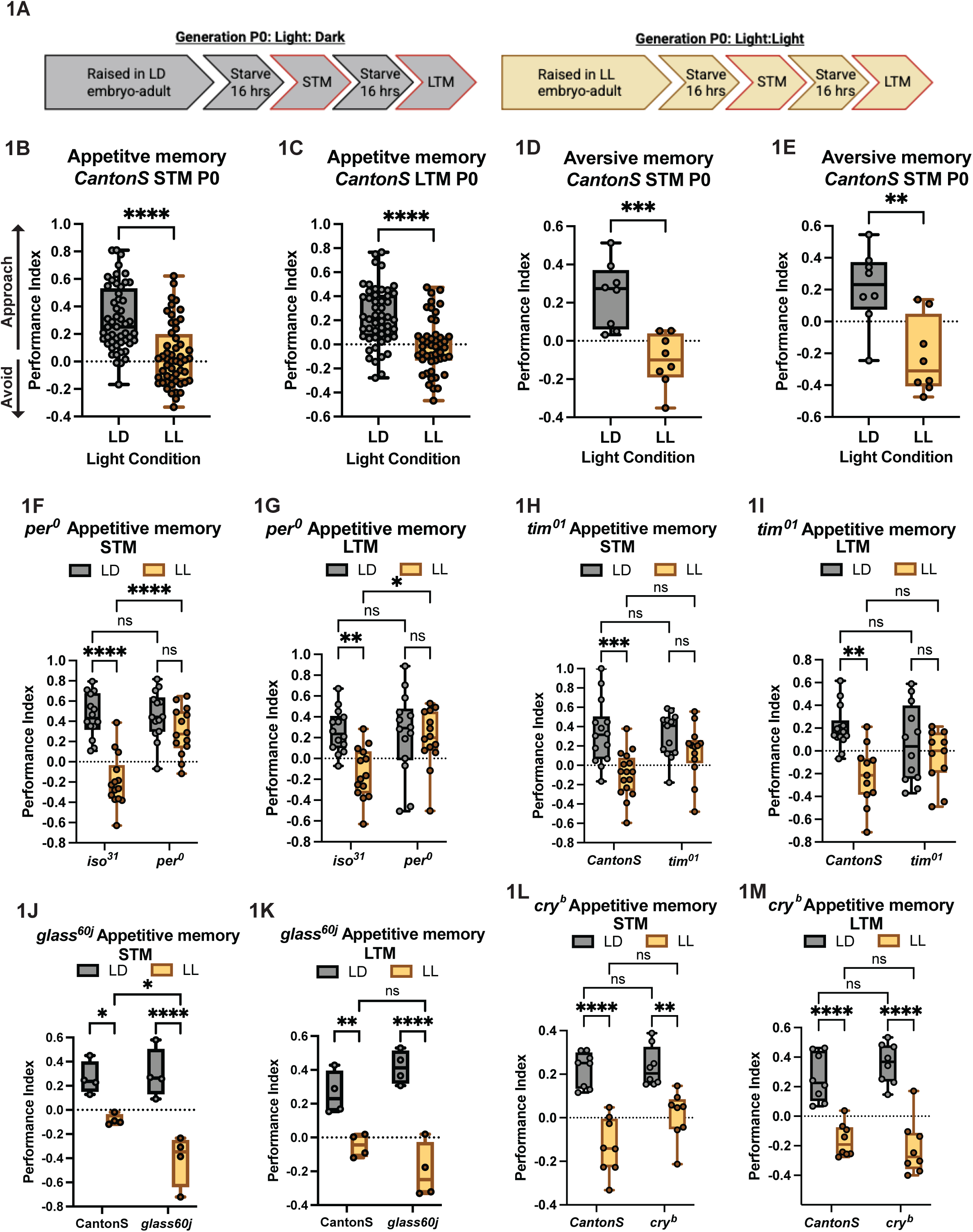
Constant light impairs associative memory in a clock-gene dependent manner. (A) Schematic of associative memory protocol. Performance Index (PI) = (# of flies choosing sugar associated odor - # of flies choosing water associated odor)/(total number of flies). Performance Index = (PI 1 + PI 2)/(2). (B and C) CantonS red eyed adult flies raised in constant light have impaired appetitive short-term (B) and long-term (C) memory performance; n = 10 biological replicates, 52 replicate assays. (D and E) CantonS red eyed flies raised in constant light have impaired aversive short-term (D) and long-term (E) memory performance; n = 3 biological replicates, 12 replicate assays. (F and G) The *per clock gene* is required for impaired short-term (F) and long-term (G) appetitive memory in constant light; n = 2 biological replicates, 14 replicate assays. (H and I) short-term memory is impaired in *tim^01^* mutants in LD and LL and long-term appetitive memory is resistant to the effect of constant light; n = 3 biological replicates, 14 replicate assays. (J and K) Photoreceptor cells (*glass^60j^*) are not required for appetitive short-term (J) or long-term (K) memory in constant light; n = 1 biological replicate, 4 replicate assays. (L and M) Cryptochrome (*cry^b^*) mutants perform short-term (L) and short-term (M) appetitive memory like wild-type flies in constant light; n = 2 biological replicates, 8 replicate assays. One-way ANOVA, Tukey’s multiple comparison test, mean ± SEM. Each dot represents at least 50 flies. *p ≤ 0.05, **p ≤ 0.01, ***p ≤ 0.001, ****p ≤ 0.00001, ns, not significant.

We next asked if this behavior was generalizable to other deviations in standard light cycles and consistent across genotypes and odors. We found that memory was likewise impaired if flies were exposed to a jetlag paradigm (9-hour phase advance of light every 48 hours, Figure S1F-G) throughout life. Also, this impairment was not restricted to red eyed flies, as *iso^31^* white-eyed flies were similarly affected by exposure to constant light (Figure S1H-I). Additionally, appetitive memory using another set of odors (IAA/EA) produced the same effect (Figure S1J-K), suggesting that this phenomenon is not restricted to a specific light disruption protocol, genotype or odor.

Constant light exposure can impact physiology. For instance, it is possible that constant light exposure alters the flies’ ability to sense sugar, which could explain the results presented thus far. However, we found that flies exposed to constant light, like flies exposed to LD, can sense and display chemotaxis towards increasing sugar concentrations (Figure S1L). Additionally, LD exposed flies can associate varying concentrations of sugar with an odor and properly execute both short- and long-term associative memory. However, flies exposed to LL do not display memory consolidation with any concentration of sugar paired with an odor (Figure S1M-N).

### Circadian clock genes, but not photoreceptor cells, are required for memory impairment under constant light conditions

Drosophila display time-of-day differences in memory consolidation, implicating the circadian clock in this regulation (Barrio-Alonso et al., 2023; Fropf et al., 2014). As constant light disrupts clock function, we asked if the effect of constant light on memory was mediated by the molecular clock. Although exposure to constant light impaired associative memory in wild-type flies, it did not do so in *per^0^* mutants (Benzer, 1971) (Figure 1F-G). Likewise, short-term memory (STM) was unaffected by light in *tim^01^* mutants (Sehgal et al., 1994). Although long-term memory (LTM) of *tim^01^* also showed no effect of LL, it was even impaired in LD (Figure 1H-I). These results suggest that at least some clock genes are required for memory impairment under constant light conditions.

In flies, either the visual system or a dedicated circadian photoreceptor, Cryptochrome (CRY), can entrain the clock to light input (Agrawal et al., 2017; Rieger et al., 2003; Stanewsky et al., 1998; Wheeler et al., 1993). Although the visual system is not essential, the sensitivity of genetically eyeless, opsin-depleted, or blind flies to light-mediated synchronization of behavioral rhythmicity is much lower than normal (Stanewsky et al., 1998). Mutants of the *glass* (*gl*) gene provide a good model to test the role of the visual system as these mutants lack normal development of photoreceptor cells, but maintain normal circadian rhythms (Moses et al., 1989; Vosshall and Young, 1995). We found that like wild-type flies, *gl*^60^ mutants have reduced appetitive memory performance in constant light (Figure 1J-K). Similarly, we found that mutations in *cry* (*cry^b^*) have impaired appetitive memory in constant light (Figure 1L-M). We cannot exclude the possibility that CRY and the visual system can substitute for each other in mediating effects of constant light on memory, but each one alone is dispensable.

### Constant light-induced changes in memory formation are intergenerationally inherited

Adverse conditions encountered by individuals can sometimes cause deficits that are transmitted to the next generation (Perez and Lehner, 2019). We wondered if the memory impairment seen in flies exposed to constant light could be intergenerationally inherited by progeny returned to a standard LD cycle. We trained and tested either LD or LL parents (P0), as described above, to confirm changes in memory. Flies exposed to constant light did not form an associative memory. To test whether progeny of flies exposed to constant light inherit impaired memory, we allowed naive parents to mate, and lay eggs for 36 hours in LD or LL. Subsequently embryos from constant light were moved to LD, while a subset of embryos were maintained in LL or in LD from one generation to the next and served as additional controls. We then performed appetitive memory training and testing on F1 adults (Figure 2A). We found that unlike LD controls, but like LL controls, progeny from LL parents that were moved to LD (LL-LD) to develop and eclose lacked short- and long-term appetitive memory (Figure 2B-C). We next asked if this inheritance could be generalized to other types of light disruption or genetic background. We found that like parental learning and memory, intergenerational inheritance of this behavior also occurred in *iso^31^* wild-type flies (Figure S2A-B) and with a chronic jetlag paradigm (Figure S2C-D).

**Figure 2:**
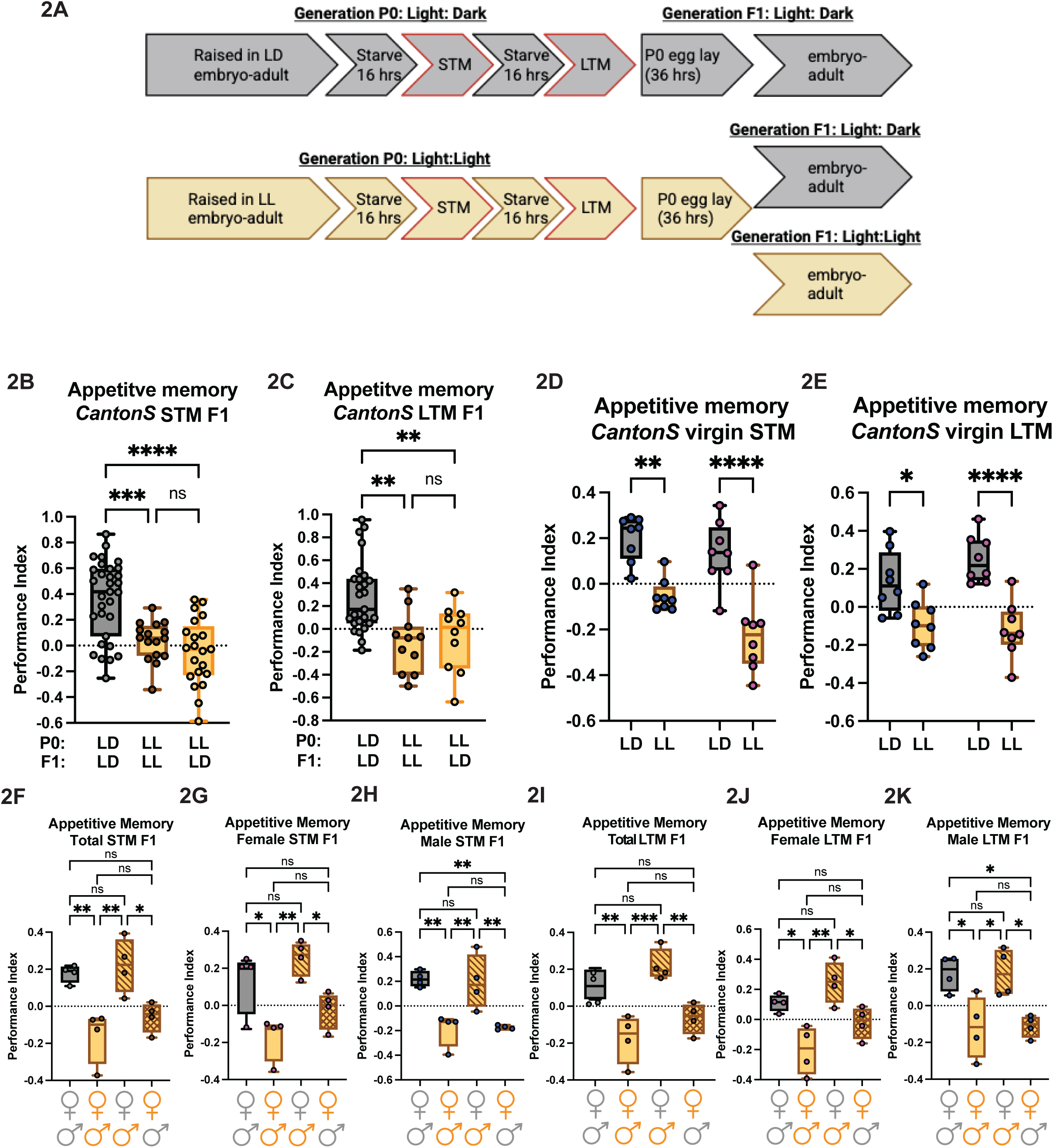
Impaired appetitive memory is intergenerationally inherited through the female germline (A) Schematic of intergenerational appetitive memory protocol. (B-C) F1 progeny of LL exposed parents have impaired short-term (B) and long-term (C) appetitive memory; n = 3 biological replicates, 10 replicate assays. (D-E) CantonS virgin males (blue dots) and females (pink dots) raised in LD perform standard appetitive short-term (D) and long-term (E), while LL-virgins are impaired; n = 2 biological replicates, 8 replicate assays. (F-K) Progeny of LD-trained mated parents (gray, LD males x gray, LD females) approach sugar rewards during short-term (F-H) and long-term (I-K) memory. Progeny of LL-trained mated parents (orange, LL males x orange, LD females) do not approach sugar rewards during short-term (F-H) and long-term (I-K) memory. Progeny of crosses where only females were exposed to LL show impaired short-term (F-H) and long-term (I-K) memory, while progeny of crosses where only males were exposed to LL have normal short-term (F-H) and long-term (I-K) memory; n = 2 biological replicates, 8 replicate assays. One-way ANOVA, Tukey’s multiple comparison test, mean ± SEM. Each dot represents at least 50 flies. *p ≤ 0.05, **p ≤ 0.01, ***p ≤ 0.001, ****p ≤ 0.00001, ns, not significant.

To investigate the basis of the intergenerationally-inherited impaired memory, we asked if there were any obvious changes in reproductive fitness or development of the LL-exposed flies. We determined reproductive ability, including counting the number of progeny, from parents exposed to constant light or LD. We found no appreciable differences in the length of the reproductive span (i.e. number of days flies lay fertilized eggs) or the total number of progeny eclosed (Figure S3A-B). These results suggest that reproduction is not affected by deviations in the standard light cycle and likely does not contribute to the observed behavioral changes. Flies exposed to constant light eclose a day earlier than flies exposed to LD (Figure S3C), but other than this and reported effects of LL on the circadian rhythm of eclosion (Skopik and Pittendrigh, 1967), no developmental deficits were observed.

### Female germlines transmit LL-induced memory impairment

Intergenerational inheritance can be transmitted through the male or female germline, depending on the behavior being investigated (Brennecke et al., 2008; Chen et al., 2016a; Dias and Ressler, 2014; Gapp et al., 2014; Hibshman et al., 2016; Moore et al., 2019; Perez et al., 2017; Stern et al., 2014; Valtonen et al., 2012; Wei et al., 2014a). To determine the contributions of the male and female germlines in transmission of this intergenerational behavior, we asked whether sperm or oocytes from trained parents (P0) could induce the behavior in the next generation (F1). First, we took advantage of the fact that we can collect virgin males and females and train and test them for memory and then mate them in specific combinations. Following exposure to LD or LL, we trained and tested virgin males and females for appetitive short- and long-term memory. LD exposed virgin males and females showed normal short- and long-term memory (Figure 2D-E). Like mated flies exposed to constant light, virgin male and female wild-type flies maintained in LL did not display short- or long-term memory (Figure 2D-E). We then mated memory trained virgin males (LL) or females (LL) to untrained virgin females (LD) or males (LD) respectively, in LD for 36 hours (control crosses of LD males x LD females, or LL males to LL females were maintained in LD or LL, respectively). After mating, we removed P0 parents and allowed F1 progeny to develop and eclose. We then performed appetitive or aversive memory assays on F1 flies. We found that F1 progeny of virgin females exposed to constant light inherited impairments in their ability to consolidate appetitive and aversive memory (Figure 2F, I, Figure S3D, 3G), while progeny of virgin males exposed to constant light had standard memory performance (Figure 2F, I, Figure S3D, 3G). Impairments in appetitive or aversive memory were inherited by both male and female F1 progeny (Figure 2G-H, 2J-K, Figure S2E-F, S2H-I), if mothers were exposed to constant light. These results suggest a role for oocytes specifically in intergenerational inheritance of LL-induced deficits in memory.

### The germline is required for associative memory

The germline is essential for the propagation of future generations and this transgenerational effect on memory is transmitted via females. This raises the question of whether the female germline itself affects memory. The germline is essential for aversive memory in *C. elegans* (Kaletsky et al., 2020), however, the role of the germline in appetitive associative memory in Drosophila has not been investigated.

Overexpression of *bam (bag of marbles)* eliminates oogenic stem cells, while somatic stem cell populations and male germline cells are unaffected (Ohlstein and McKearin, 1997). Therefore, we used this tool to explicitly investigate the role of the female germline in learning and memory in LD. Using two broad germline drivers, nanos-Gal4 and GAL4::VP16-nanos (a stronger nanos driver) to drive *UAS-Bam*, we screened for germline involvement in parental learning and memory. We found that eliminating the ovarian stem cell niche suppressed flies’ ability to perform standard appetitive memory (Figure S3J-M). To narrow down the specific cell population within the germline that is required for memory, we used more specific Gal4 drivers. Expressing *Bam* in follicle cells using tj-Gal4 (Figure S3N-O), or in precursor cells in the embryonic epidermis, the stalk cells, and follicle cells at the anterior and posterior poles of the egg chamber using 7021-Gal4, also suppressed typical appetitive memory performance (Figure S3P-Q). Expressing *UAS Bam* in the embryonic dorsal midline, PNS, larval ventral nerve cord, segmentally in the nervous system, and in stalk and follicle cells using 7023-Gal4 did not affect memory performance (Figure S3R-S). Together these results suggest that a functional germline is required for appetitive memory.

### LL-induced impaired memory is transmitted through three generations

The data above indicate that deviations in standard LD cycles induce behavioral changes that are transmitted to the next generation. However, during training, fertilized eggs could be exposed to constant light while still inside the mother, so F1 behavior cannot be considered truly transgenerational. True transgenerational epigenetic inheritance persists beyond the first generation, and the duration of this persistence often depends on the specific phenotype; therefore, we tested how many generations this change in memory persists once progeny of LL parents are moved to LD. We exposed CantonS red eyed or *iso^31^* parents (P0) to LD or to constant light and performed associative memory experiments to confirm impaired memory. We next moved egg lays from a subset of LL parents to LD and performed short- and long-term memory experiments in progeny (control LL and LD populations were maintained in their respective light cycles for subsequent generational testing). We found that parental (P0) constant light exposure induces short- and long-term behavioral changes in progeny moved to LD in three subsequent generations, F1-F3 (Figure 3, Figure S2A-B). Fourth generation (F4) descendants properly associate a CS+ pairing, resuming standard trained behavior (Figure 3, Figure S2A-B). Similarly, exposing CantonS red eyed parents to a chronic jetlag protocol and subsequently moving progeny to a standard LD light cycle also results in three generations of deficits in memory before returning to baseline in the F4 (Figure S3C-D). Interestingly, the performance index of F1-F3 LL-LD progeny does not change until memory is rescued in the F4 generation.

**Figure 3:**
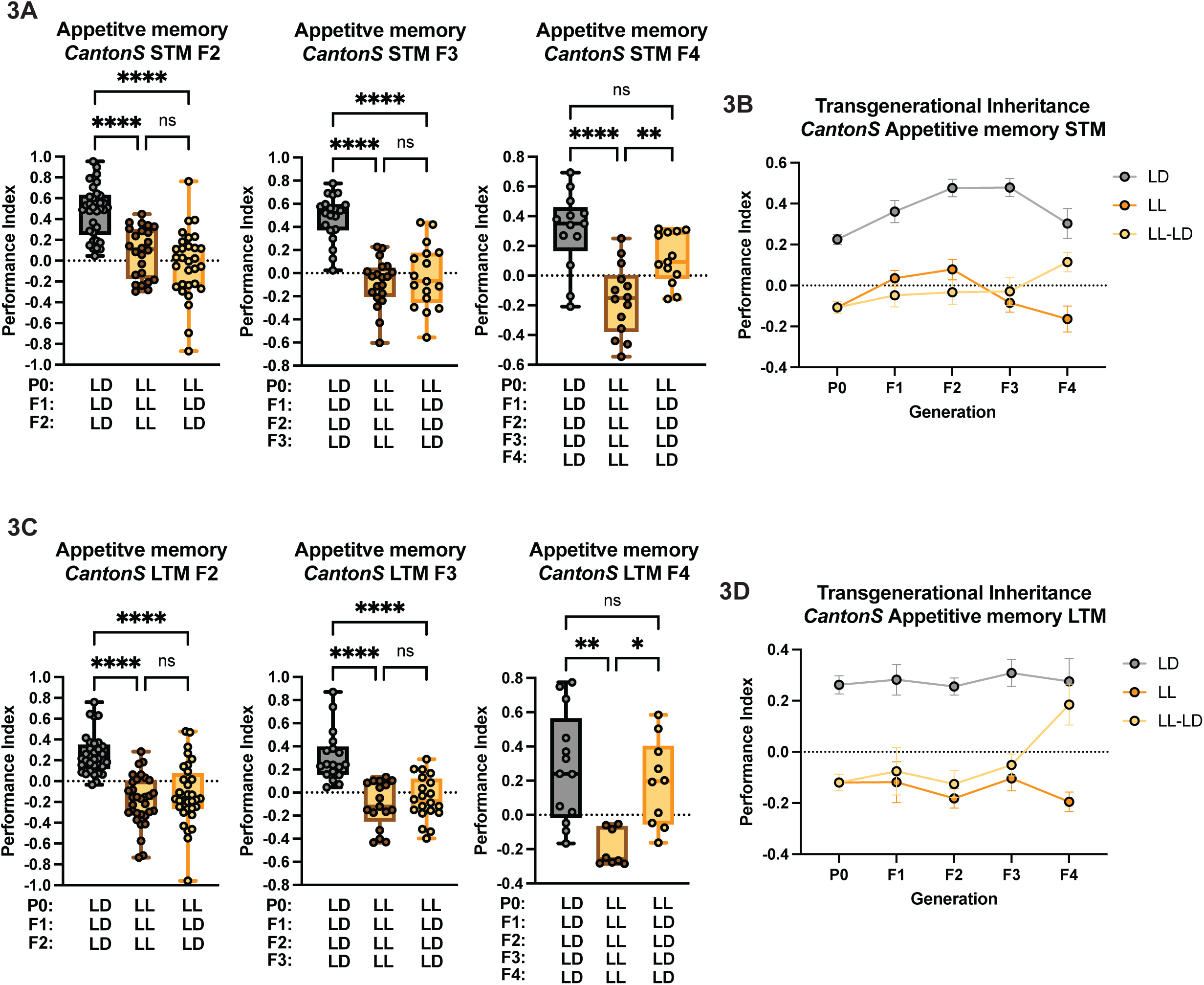
LL-induced impairments in memory are inherited for three generations in progeny raised in LD (A-D) Progeny of LL-parents raised in LD inherit impaired short-term memory (A, B) and long-term memory (C, D) from generation F1-F3, before returning to standard appetitive memory responses in the fourth generation. One-way ANOVA, Tukey’s multiple comparison test, mean ± SEM. n = ≥ 2 biological replicates, ≥ 12 replicate assays. Each dot represents at least 50 flies. *p ≤ 0.05, **p ≤ 0.01, ****p ≤ 0.0001, ns, not significant.

### The microRNA and the endogenous small interfering RNA pathway are not required for appetitive memory

To identify possible mediators of this transgenerational behavior, we investigated the role of candidate TEI regulators that have been previously implicated in regulating transmission across generations; these include microRNAs, endogenous small interfering RNAs (endo-siRNAs), and piwi RNAs (piRNAs) (Fitz-James and Cavalli, 2022; Perez and Lehner, 2019). We used RNAi to knockdown expression of specific pathways with five drivers, including Actin-Gal4 (full body), nanos-Gal4 (germline), nSyb-Gal4 (pan-neuronal), 201y-Gal4 (α/β/γ mushroom body) (Crocker et al., 2010), and 16A06-Gal4 (broad mushroom body), and raised all flies in LD. We focused on the germline and brain because the germline is required for associative memory and generation of future progeny, and memory is executed in the brain, specifically in the mushroom body (De Belle and Heisenberg, 1994; Dubnau et al., 2001; Erber et al., 1980; Heisenberg et al., 1985; McGuire et al., 2001; Mizunami et al., 1998). We reasoned that if knockdown of specific small RNA pathways in LD affected appetitive memory in normal light:dark cycles, then those pathways were candidates for mediating effects of constant light.

Both the endo-siRNA and miRNA pathways are crucial for gene regulation, with endo-siRNAs derived from endogenous RNA precursors, and miRNAs from stem-loop RNA precursors. Endo-siRNAs and miRNAs are processed by Dicer and stabilized by argonaute proteins. We found that knockdown of Dicer-1 (Dcr-1), an endoribonuclease which functions in miRNA and endo-siRNA gene silencing, impaired long-term memory when knocked down in the full body or pan neuronally; knockdown in the mushroom body disrupted STM and LTM (Figure 4A, Figure S4A). However, neuronal knockdown of Dicer-2 (Dcr-2), a member of the RNase III family of double-stranded RNA specific endonucleases that cuts dsRNA into endo-siRNAs for the RNAi pathway, yielded inconsistent results with effects on STM but not LTM (Figure 4A, Figure S4B). Additionally, knockdown of Argonaute-1 (AGO1), an argonaute/Piwi family protein, which interacts with microRNAs to form miRNA-induced silencing complexes, or Argonaute-2, an argonaute/Piwi family protein that interacts with siRNAs to form RNA-induced silencing complexes, using any driver did not affect short- or long-term memory (Figure 4A, Figure S4C, S5A). Overall, knockdown of the miRNA and endo-siRNA pathways does not yield consistent effects on memory, suggesting that these pathways are not required for appetitive memory execution, and so are unlikely to mediate effects of LL on memory.

**Figure 4:**
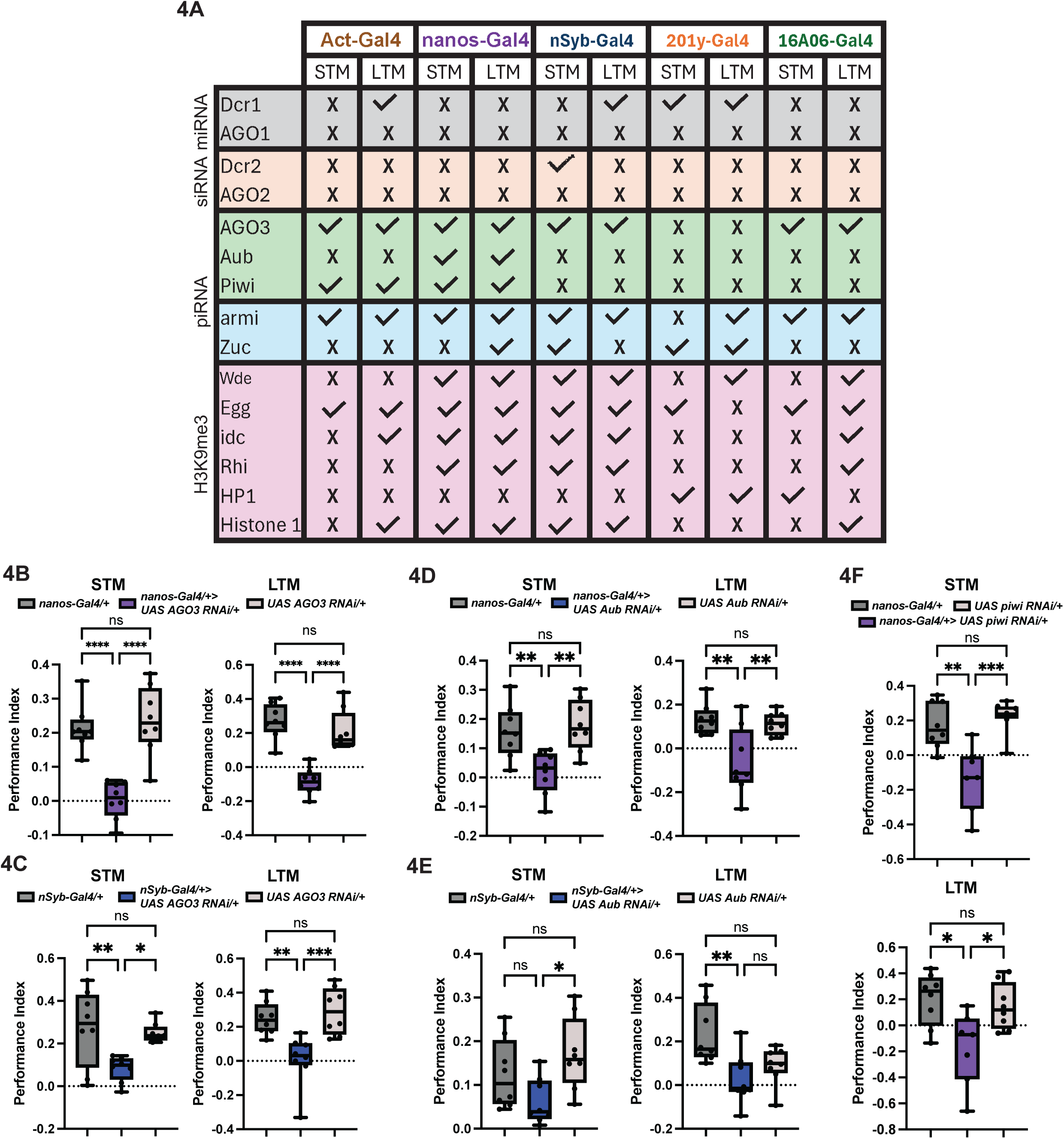
The piRNA pathway, but not the miRNA or endo-siRNA pathway, is required for associative memory. (A) The miRNA pathway (gray, Dcr-1 and AGO1) and the endo-siRNA (orange, Dcr-2 and AGO2) are not consistently required for appetitive memory in LD. The PIWI clade argonautes (green, AGO3, Aub, and Piwi) and downstream proteins are consistently required both primary and ping-pong/secondary piRNA biogenesis (blue, armi and Zuc) are required for STM and LTM. Only piRNA pathway H3K9me3 function (pink, Wde, Egg, idc, Rhi, HP1, and Histone 1) are required for STM and LTM associative memory in LD. (B-C) AGO3 is required in the germline (B) and in neurons (C) for STM (left) and LTM (right). (D-E). Knockdown of Aub in the germline (D) or in neurons (E) impairs appetitive STM (left) and LTM (right). (F) *nanos-Gal4>UAS piwi RNAi* impairs STM (top) and LTM (bottom) in LD. n = 2 biological replicates, 8 replicate assays. One-way ANOVA, Tukey’s multiple comparison test, mean ± SEM. Each dot represents at least 50 flies. *p ≤ 0.05, **p ≤ 0.01, ***p ≤ 0.001, ****p ≤ 0.00001, ns, not significant.

### The piRNA pathway is required for appetitive memory

The PIWI-interacting RNA pathway (piRNA) is a conserved system, is expressed in the nervous system, and has been implicated in learning and memory and transgenerational behaviors (Grentzinger et al., 2012; Kaletsky et al., 2020; Moore et al., 2019; Rajasethupathy et al., 2012). We found that knockdown of DCR1 affected appetitive memory, while knockdown of DCR2 has no effect (Figure 4A, Figure S4A-B). DCR1, but not DCR2, interacts with the piRNA pathway (Megosh et al., 2006) so we reasoned that the piRNA pathway represented a strong candidate to regulate learning and memory and its inheritance. Using an *UAS-RNAi* knockdown approach we found tissue specific requirements for the piRNA pathway for appropriate learning and memory.

The piRNA pathway is regulated via three argonautes. We found that all three PIWI clade argonautes — P element-induced wimpy testis (Piwi), Aubergine (Aub), and Argonaute 3 (AGO3) — were required in the germline (nanos-Gal4) for STM and LTM (Figure 4A-B, D, F, Figure S5B-D), while AGO3 and Aub were additionally required pan neuronally for STM and LTM (Figure 4A-C). Not surprisingly, the full body knockdown (Act-Gal4) of *piwi* also produced STM and LTM phenotypes (Figure 4A, Figure S5D). Because we found a role for all three argonautes we tested other downstream molecular components of the piRNA pathway that function in primary piRNA biogenesis in both somatic and germline cells (Piwi) or only in the germline-restricted secondary/ping-pong pathway (Aub and AGO3). Following their transcription, piRNA intermediates are cleaved at their 5’ end by the mitochondria-bound endonuclease, Zucchini (Zuc). Loss of Zuc results in a dramatic reduction of piRNAs (Malone et al., 2009) and causes delocalization of Piwi, but not Aub or AGO3, suggesting Zuc aids Piwi in the piRNA loading process (Olivieri et al., 2012). Loss of Zuc impaired STM and LTM when knocked down in the germline and when knocked down in the mushroom body (Figure 4A, Figure S6A). We also knocked down Armitage (Armi), an RNA helicase. Although little is known about its mechanism of action, mutation of Armi disrupts localization of Piwi, and levels of piRNAs decrease dramatically (Olivieri et al., 2012). We found that Armi was required broadly for STM and LTM. RNAi knockdown of Armi in the full body, in the germline, pan neuronally or in the mushroom body impaired memory (Figure 4A, Figure S6B). Together, these data suggest that both the primary and secondary piRNA biogenesis pathways are required broadly in the germline and brain for associative learning.

### The H3K9me3 function of piRNAs is required for learning and memory

The PIWI pathway regulates transposons, maintains germline stem cells, and mediates epigenetic modification in germline and somatic cells. Immediately upon fertilization piRNAs are deposited into the developing embryos as maternal piRNAs. *piwi^1^/piwi^2^* (Lin and Spradling, 1997) transheterozygotes and *piwi^2^* homozygous mutants in LD show impaired STM and LTM (Figure 5A). To test the role of maternal piRNAs, we mated female *piwi^1^* heterozygotes or *piwi^2^* homozygotes to wild-type control background males, and tested progeny. We found that progeny devoid of maternal piRNAs have normal memory (Figure 5B). Therefore, *piwi,* but not its maternal piRNA role, are required for appetitive memory.

**Figure 5:**
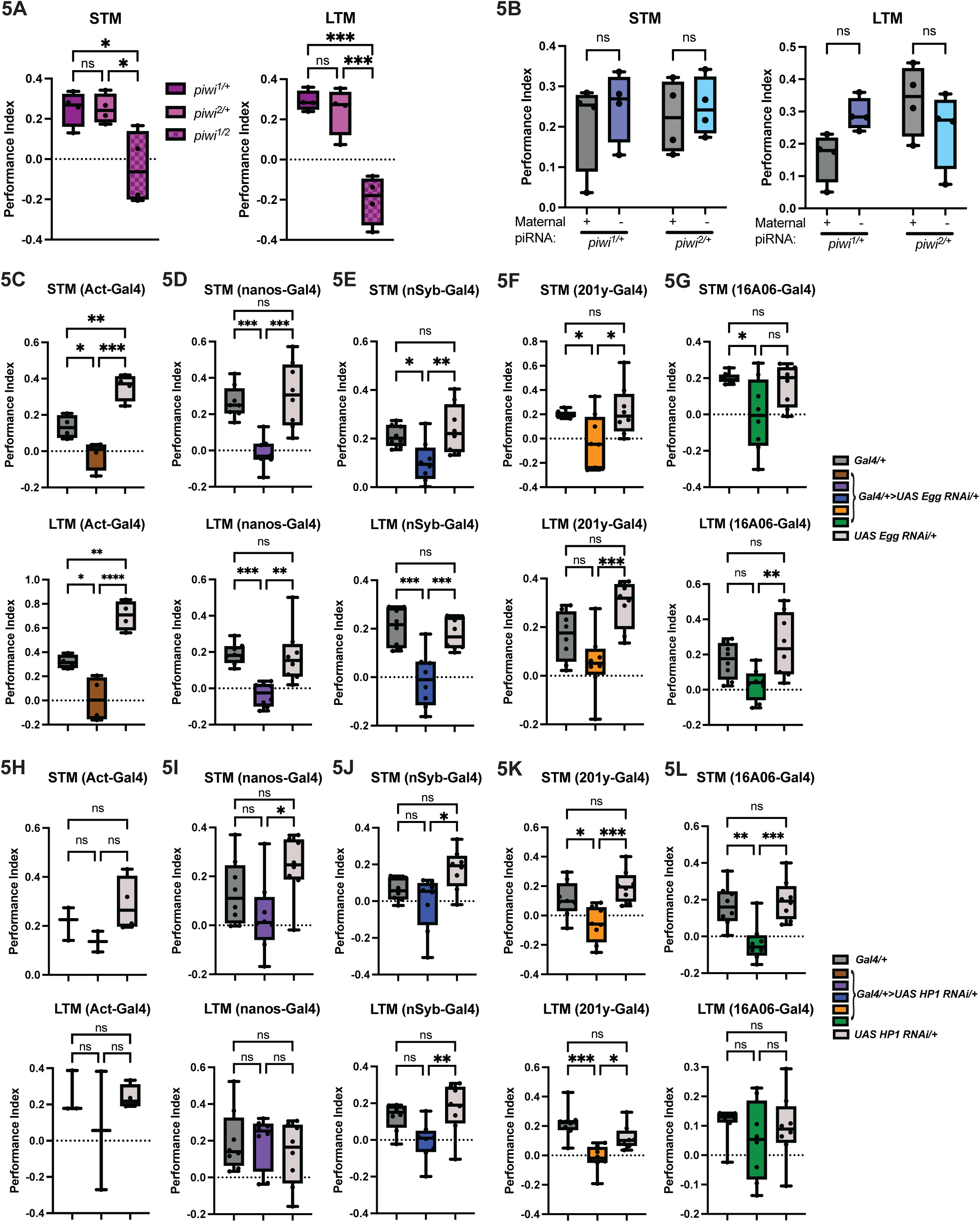
Piwi-dependent H3K9me3 deposition is required for appetitive memory (A) *piwi^1/2^* transheterozygote mutants do not display appetitive STM (left) or LTM (right) compared to heterozygote controls; n = 1 biological replicate, 4 replicate assays. (B) Regardless of the presence of maternal piRNAs there is no difference in STM (left) or LTM (right) performance; n = 1 biological replicate, 4 replicate assays. (C-G) Eggless is required full body (C), in the germline (D), in neurons (E), and in the mushroom body (F-G) for STM and LTM; n = 2 biological replicates, 8 replicate assays. (H-L) HP1 RNAi does not affect appetitive STM or LTM full body (H), in the germline (I), or in neurons (J), but impairs memory when knocked down in the mushroom body (K-L); n = 2 biological replicates, 8 replicate assays. One-way ANOVA, Tukey’s multiple comparison test, mean ± SEM. Each dot represents at least 50 flies. *p ≤ 0.05, **p ≤ 0.01, ***p ≤ 0.001, ****p ≤ 0.00001, ns, not significant.

Piwi proteins, along with cofactors and piRNAs, silence targets, including transposons and mRNAs, by guiding the deposition of repressive heterochromatin Histone 3 Lysine trimethyl marks (H3K9me3) via two mechanisms. For one, deposition of H3K9me3 can be accomplished by the piRNA-RISC complex via Eggless (Egg, dSETDB1), a histone methyltransferase, and its cofactor Windei (Wde, mAM/MCAF1 (Koch et al., 2009)). Mutations in Egg or Wde lead to decreases in H3K9me3 deposition in both germ and somatic cells, resulting in a reduction of mature piRNAs (Koch et al., 2009; Rangan et al., 2011). We found that knockdown of either Egg or Wde affected STM and LTM (Figure 5C, S7A). Knockdown of either Egg or Wde in the germline or in neurons impaired both STM and LTM (Figure 4A, 5D-E, Figure S7A). Knockdown of Egg in the whole body or in the mushroom body also abrogated STM and LTM (Figure 4A, 5F-G).

A second mechanism by which piRNAs can direct H3K9me3 silencing is via DNA binding proteins. identity crisis (idc) is a Kruppel box-associated zinc finger (KRAB-ZNF) binding protein, which is involved in H3K9me3 mediated transcriptional silencing of repetitive DNA elements (Shapiro-Kulnane et al., 2022). Knockdown of *idc* in the germline or in neurons impaired STM and LTM, while knockdown in the full body or using the broad mushroom body driver (16A06-Gal4) only affected LTM (Figure 4A, Figure S7B).

The precise mechanism by which Piwi proteins influence chromatin remains elusive; however once H3K9me3 marks are deposited, Heterochromatin Protein 1 (HP1) and its homolog, Rhino (Rhi, transposon silencing specific) are thought to bind H3K9me3 (Bannister et al., 2001; Lachner et al., 2001) and interact with Piwi proteins (Brower-Toland et al., 2007) suggesting that HP1 directs Piwi proteins and its’ bound piRNAs to execute silencing. Interestingly, HP1 and Rhi have tissue specific requirements for memory. Knockdown of HP1 did not affect memory when knocked down full body or in the germline but eliminated STM and LTM when knocked down in the mushroom body (Figure 5H-L), while Rhi was required in the germline and in neurons for appetitive memory (Figure 4A, Figure S7C). Finally, we found that Histone1 was required pan neuronally for STM and LTM; knockdown with a broad mushroom body driver impaired LTM (Figure 4A, Figure S8A). Due to the multitude of evidence surrounding the H3K9me3 role of Piwi argonautes, we speculate that this function of Piwi is essential for appetitive memory.

### Transgenerational effects of constant light are mediated by the PIWI pathway

PIWI subfamily proteins are expressed and easily visualized in the germline where they are required for germ stem cell maintenance. We have found that transmission of LL-induced deficits in appetitive memory occurs via the female germline and that the piRNA pathway is required in the female germline for memory. Based on this, we speculated that female germline PIWI expression might be affected by LL. We exposed flies expressing *EGFP::Piwi* (Sienski et al., 2012) to either LD or LL for three generations. Constant light increased nuclear Piwi expression compared to LD (Figure 6A-C). We then asked whether high PIWI expression was inherited by LD-raised progeny of mothers exposed to constant light. Naïve F1 progeny of P0 mothers exposed to constant light maintained a similar level of nuclear *EGFP::Piwi* expression in the ovary as compared to their LL-mothers (Figure 6B-E). Next, we asked how many generations the increased expression of Piwi in the ovary persists after exposure to constant light. Piwi remained elevated in the germline of progeny of constant light exposed mothers for three generations before returning to basal levels in the fourth generation (Figure 6F-I), mirroring the trajectory of the inherited behavior (Figure 3).

**Figure 6:**
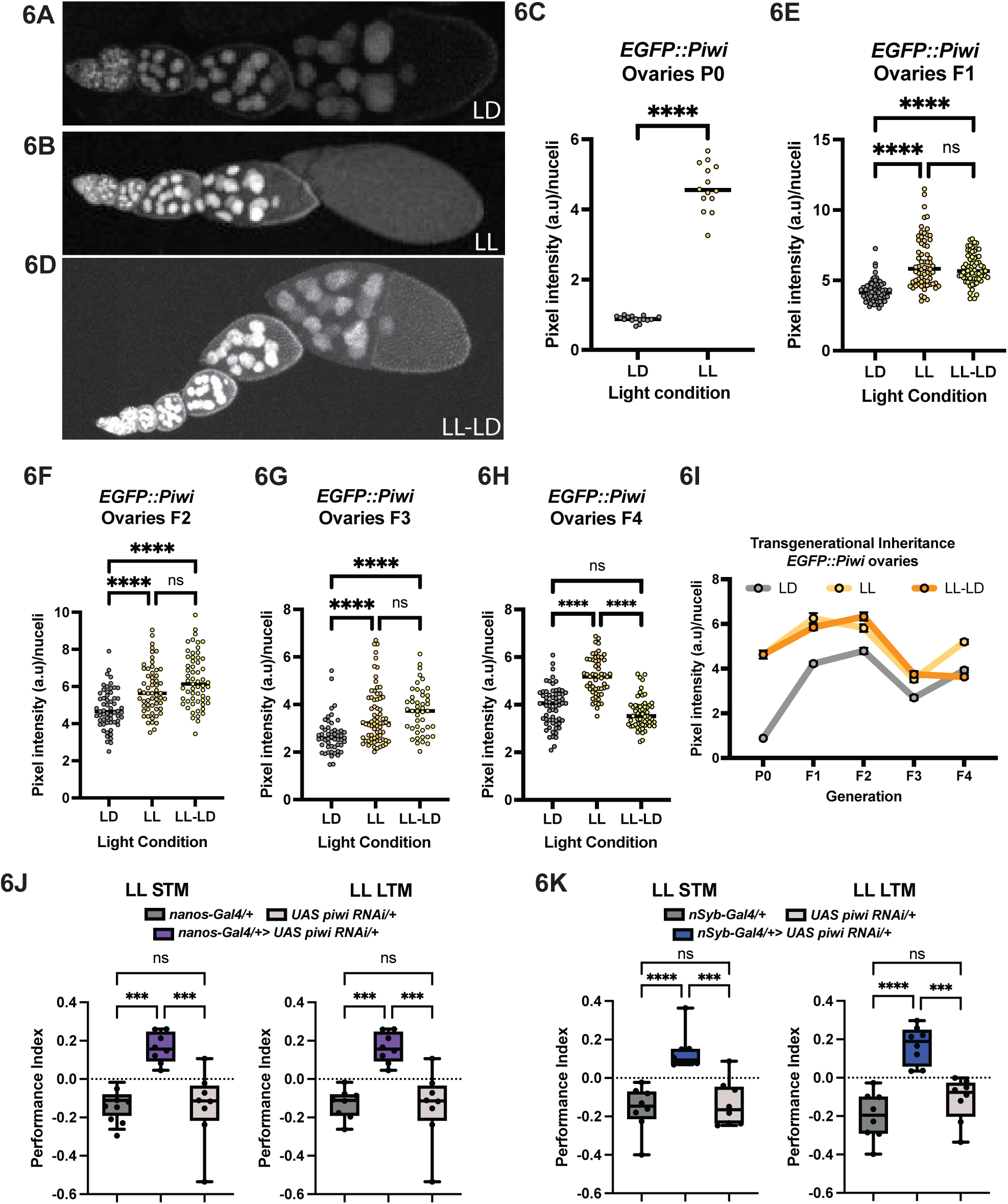
Constant light exposure regulates PIWI expression transgenerationally (A) EGFP*::Piwi* is expressed in germline nuclei in LD. (A-C) Constant light (LL) exposure, as compared to LD, increases *EGFP::Piwi* expression in germline nuclei compared to LD. Student’s t-test, mean ± SEM. (D-E) F1 Progeny of LL parents, raised in LD, have increased *EGFP::Piwi* expression in germline nuclei compared to progeny of LD parents. (F-I) Parental constant light exposure increases *EGFP::Piwi* expression in germline nuclei of F2 (F) and F3 (G) progeny raised in light:dark for three generations before returning to LD levels in the fourth (F4, H-I). (J) *nanos-Gal4>UAS piwi RNAi* restores STM and LTM in LL. (K) Reduction of *Piwi* in neurons (nSyb-Gal4) restores appetitive STM and LTM in LL. One-way ANOVA, Tukey’s multiple comparison test, mean ± SEM. For images: Each dot represents one glomeruli lobe. n = 3 biological replicates, 8-10 ovaries per condition per replicate. *p ≤ 0.05, **p ≤ 0.01, ***p ≤ 0.001, ****p ≤ 0.00001, ns, not significant. For memory assays: One-way ANOVA, Tukey’s multiple comparison test, mean ± SEM. Each dot represents at least 50 flies. n = 2 biological replicates, 8 replicate assays.

We reasoned that elevated levels of Piwi could be as deleterious to memory as reduced levels and so could underlie impaired memory in LL. As knockdown of *piwi* in the brain or the germline was sufficient to impair memory in LD, and PIWI is elevated in the germlines of LL exposed females, we knocked down *piwi* in both tissues and performed appetitive memory experiments. Knockdown of *piwi* in the germline (Figure 6J) or in neurons (Figure 6K) of flies raised in LL restored STM and LTM. These results indicate that the overexpression of Piwi is responsible for memory deficits in LL and knocking down its expression can restore appetitive memory performance.

### PIWI mediates effects of constant light on memory by modulating dopamine signaling

In addition to the germline, we found that the piRNA pathway is required in the brain, in particular in the mushroom body, for associative memory so we focused our attention on the mushroom body. The mushroom body (MB) neuropil, consisting of alpha, beta, and gamma lobes, is a compartmentalized higher brain structure that is necessary for most forms of associative memory in flies (Heisenberg, 2003; Keene and Waddell, 2007; McGuire et al., 2001). We found no appreciable morphological changes in whole brains (*nSyb-GAL4>UAS-mcd8-GFP*) or in the mushroom body (*R35B12>UAS-mcd8-GFP* and *R26E01>UAS-mcd8-GFP*) in LL flies (Figure S8B), leading us to postulate that neural transmission could be responsible for the behavior phenotype we observe.

Excessive light exposure decreases dopamine expressing neurons in rats (Romeo et al., 2013), and dopamine itself is light sensitive (Hirsh et al., 2010). Additionally, Handler et al (Handler et al., 2019) demonstrated that the Dopamine Receptor 1R1 (dA1 or dumb) and Dopamine Receptor 1R2 (damb) activity oppose each other in mediating approach vs avoidance behavior following memory training. We considered the possibility that the memory impairment in LL was caused by disrupted regulation of approach/avoidance, perhaps by changes in dopamine signaling. To determine whether dopamine signaling was affected in our LL paradigm, we exposed P0 *lexAop tdTomato.nls>UAS-Dop1R1-Tango* flies to LD or LL for one generation, dissected, and imaged brains. We found that flies raised in LL had fewer Dop1R1 puncta compared to flies raised in LD (Figure 7A-C), although puncta size was unaffected (Figure 7A-B, D). Next, we asked if Dop1R1 expression changed after memory training. We exposed *lexAop tdTomato.nls>UAS-Dop1R1-Tango* flies to LD or LL for one generation, performed memory training, dissected brains, and imaged flies. We found that flies raised in LD increased the number and size of Dop1R1 puncta after STM while flies raised in LL did not (Figure 7E-H). These data suggest that irregular light exposure dampens dopamine signaling via Dop1R1 at baseline and prevents its increase during STM.

**Figure 7:**
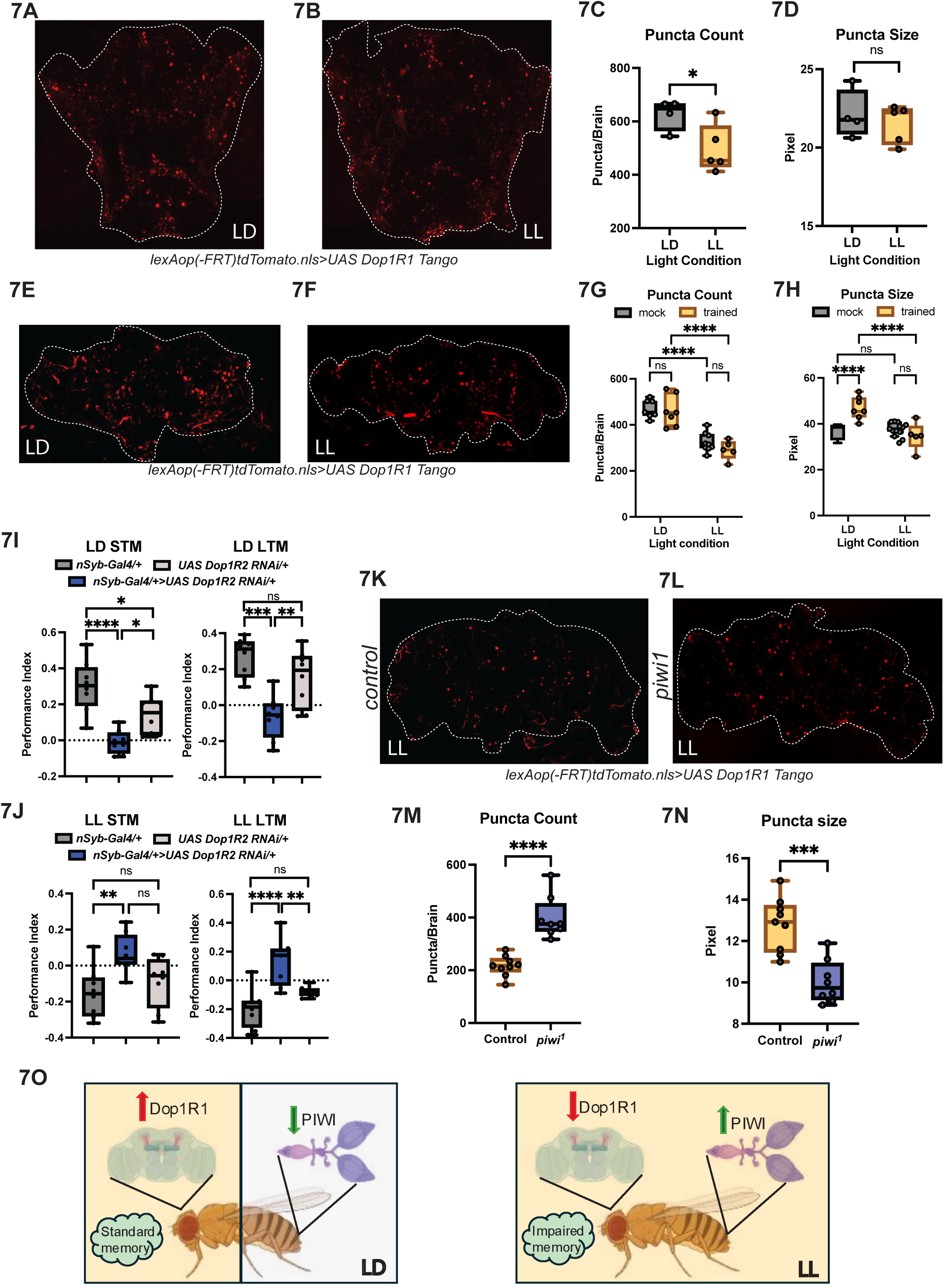
Constant light suppresses Dop1R1 activity via Piwi (A) *lexAop(-FRT)tdTomato.nls>UAS Tango Dop1R1* is expressed ubiquitously in the brain in LD. (B-D) Constant light decreases the number of Dop1R1 puncta (C) without affecting puncta size (D). (E-F) Dop1R1 expression following STM in LD (E) and LL (F). (G) STM does not alter the number of Dop1R1 puncta in flies raised in LD or LL. (H) STM increases Dop1R1 puncta size in LD, but no change is seen in flies raised in LL. (I) Knockdown of Dop1R2 in neurons impairs STM (left) or LTM (right) in LD. (J) Knockdown of Dop1R2 in neurons restores STM (left) or LTM (right) in LL; n = 2 biological replicate, 8 replicate assays. (K-L) In LL *piwi*^1^ mutants have more Dop1R1 puncta compared to control flies. (N) Dop1R1 puncta are smaller in *piwi*^1^ mutants compared to controls in LL. (O) Model of impaired memory due to constant light exposure. For images: Each dot represents one glomeruli lobe. n = 2 biological replicates, 4-9 brains per condition per replicate. *p ≤ 0.05, **p ≤ 0.01, ***p ≤ 0.001, ****p ≤ 0.00001, ns, not significant. For memory assays: One-way ANOVA, Tukey’s multiple comparison test, mean ± SEM. Each dot represents at least 50 flies. n = 2 biological replicates, 8 replicate assays.

We were unable to measure expression of Dop1R2 due to unavailability of appropriate tools, but because these two receptors counter each other, we reasoned that Dop1R2 might be increased in LL. Thus, we sought to reduce Dop1R2 activity and test appetitive memory in constant light. We raised flies in LD or in LL while knocking down Dop1R2 activity in neurons (nSyb-Gal4) and performed STM and LTM. Knockdown of Dop1R2 in LD in neurons (Figure 7I) impaired short-term and long-term memory. In LL, knockdown of Dop1R2 in neurons (Figure 7J) restored normal STM and LTM. These results suggest that constant light alters Dop1R2 activity, likely increasing its’ activity so that avoidance promoted by this receptor counters the approach behavior that would normally follow appetitive memory training.

Our data suggest that in response to constant light PIWI expression is increased, and reducing its expression corrects memory. Simultaneously, Dop1R1 activity is reduced in LL, and reducing Dop1R2 rescues LL memory suggesting that LL causes an imbalance of dopamine signaling. Together, this suggests that *piwi* regulates Dop1R1 activity in response to constant light. To test this idea, we generated *piwi^1^* mutants carrying *UAS-Dop1R1-Tango* to investigate Dop1R1 expression when *piwi* expression is absent, in LD and LL. We found that in constant light *piwi* mutants had more, but smaller, Dop1R1 puncta than control flies (Figure 7M-P). Together, these results suggest that in constant light *piwi* regulates Dop1R1 expression.

## DISCUSSION

Societies live in 24-hour environments, in which light and darkness follow a diurnal pattern and allow organisms to synchronize their behavioral and physiological rhythms. Modern lifestyles, excessive workload, shift work and short night periods involve exposure to excessive artificial light, which severely affects endogenous synchrony and increases susceptibility to numerous health disorders (Potter et al., 2016; Qian and Scheer, 2016; Videnovic and Zee, 2015). Despite evidence of the negative effects of light pollution, our current understanding of it applies only to exposed individuals (i.e. intragenerational) and generally lacks mechanistic understanding. Importantly, transgenerational effects have not been examined.

In this study we investigated the transgenerational effect of constant light exposure on appetitive and aversive associative memory in *D. melanogaster*. We show that constant light impairs short- and long-term memory (Figure 1B-E), independent of any effects on brain architecture (Figure S8B) or sensing acuity (Figure S1I). This behavior requires molecular clock genes *per* and *tim*, but it does not appear to require disruption of the clock. While LL typically disrupts the clock, *cry^b^* mutants maintain rhythmic behavior and molecular clock function under these conditions (Emery et al., 2000) and *cry^b^* mutants respond to LL with impairments of memory. It is possible that LL affects circadian function in *cry* mutants in ways that have not been discovered yet, e.g. in specific tissues not required for rhythmic rest:activity. Alternatively, PER and TIM may have non-circadian roles in the effect of LL.

LL-induced memory impairment is inherited for three generations in progeny of constant light-exposed parents raised in a standard light-dark cycle. We found that the transgenerational epigenetic inheritance is transmitted through the female germline and requires the PIWI pathway. The PIWI-interacting RNA pathway is a potent regulator of transgenerational epigenetic phenomena across a variety of species (Grentzinger et al., 2012; Kaletsky et al., 2020; Moore et al., 2019; Rajasethupathy et al., 2012). While piRNA biogenesis and secondary amplification mechanisms vary, the output of the piRNA pathway converge on the regulation of gene expression, typically via silencing. In flies (Jones et al., 2016; Tindell et al., 2020; Wakisaka et al., 2019), worms (Huang et al., 2023; Kim et al., 2018; Sun et al., 2018), and mice (Huang et al., 2023; Lee et al., 2011), piRNAs are also expressed outside of the germline where they regulate stable silencing of somatic gene expression via H3K9me3 deposition. Our results present novel regulation of piRNA expression by light and demonstrate that this class of small RNAs mediates the transgenerational effect of constant light on associative memory via Dop1R1 regulation (Figure 7Q).

PIWI has been implicated in learning and memory in aplysia (Rajasethupathy et al., 2009), worms (Moore et al., 2019), and mice (Leighton et al., 2019), and it is required for transgenerational inheritance in worms (Moore et al., 2019) and flies (Casier et al., 2019; Fabry et al., 2021) although in the context of a different behavior. Our data overall indicate that regulation of piRNAs by proteins such as the argonautes AGO3, Aubergine, and Piwi, the RNA helicase Armitage, and the piRNA maturation factor Zucchini, is required in the germline and brain for appetitive memory in normal light:dark cycles. Additionally, molecular components required for H3K9me3 deposition and maintenance including, Wde, Egg, Rhino, idc, HP1, and Histone1 are also required in the germline and in neurons. Thus, we found that the PIWI pathway regulation of H3K9me3 is important for appetitive memory in a light-dark cycle.

We show that exposure to constant light or a chronic jet lag paradigm impairs associative appetitive and aversive memory in progeny raised in a standard light:dark cycle for three additional generations prior to reverting in the fourth generation (Figure 3, Figure S3). This impaired memory is accompanied by an upregulation of PIWI in the germline (Figure 6). Interestingly, the performance index in memory assays and PIWI expression remains constant during the transgenerational inheritance. So, rather than the steady loss of a direct target, which might result in a declining fraction of animals exhibiting the behavior in each generation (as in mortal germline or small RNA-triggered stable silencing of GFP transgenes (Ashe et al., 2012; Bagijn et al., 2012; Luteijn et al., 2012; Shirayama et al., 2012)) our data support an all-or-none threshold model, where behavior is consistent from P0 to F3 due to the elevated levels of Piwi, then is lost at F4 when Piwi expression returns to baseline levels.

Dopamine plays a crucial role in both learning and memory by signaling the perceived saliency of stimuli, modulating memories via synaptic plasticity, and supporting memory consolidation (Castillo Díaz et al., 2021). It was previously reported that excessive exposure to bright light can reduce the number of dopamine positive neurons and even decrease dopamine and its metabolites in the brain (Romeo et al., 2013). In *Drosophila*, Dop1R1 and Dop1R2 play opposing roles in olfactory memory regulation at the behavioral level, with Dop1R1 essential to approach following appetitive pairing while Dop1R2 is necessary for avoidance via potentiation following backward pairing, when the CS+ is presented after the US (Berry et al., 2018; Handler et al., 2019; Kim et al., 2007; Qin et al., 2012). Here, we show that pan neuronal Dop1R1 expression is dampened by constant light and knockdown of Dop1R2 in neurons restores appetitive memory. Together this suggests that constant light renders Dop1R1 activity low and Dop1R2 activity high, increasing avoidance and thereby reducing the approach response to the CS+.

Constant light dampens neuronal Dop1R1 activity, and increases germline PIWI expression transgenerationally, mirroring impaired memory performance. Given that *piwi* knockdown in the brain restores LL memory, we infer that PIWI is also elevated in the brain. Whether the LL effect on *piwi* occurs first in the brain or the germline is not known, but transmission to the next generation is clearly through the germline. We show that Dop1R1 activity can be rescued in *piwi* mutants in constant light. This suggests a model in which constant light elevates PIWI expression which causes high levels of silencing by H3K9me3 on the Dop1R1 coding region. How may PIWI regulate dopamine? First, PIWI-interacting RNA pathway members, tested in this paper, colocalize with Dop1R1 and Dop1R2 throughout the lifetime of the animal (Li et al., 2022). Furthermore, within the coding region of the Dop1R1 are two transposable elements (FBti0019373-RA, and FBti0019374-RA), representing potential silencing sites for the PIWI pathway. This would fit our model that in response to constant light PIWI expression is elevated, resulting in hypersuppression of Dop1R1 expression and hyperactivity of Dop1R2.

To address the role of different small RNA pathways in appetitive memory, we knocked down many components of these pathways in several different tissues. While this overall indicated a role for the piRNA pathway, there were some discrepant effects on STM/LTM with RNAi knockdown in different tissues. For instance, in a couple of cases knockdown in the mushroom body yielded a stronger phenotype than knockdown in the whole body. This could result from a stronger driver for the mushroom body, differences in the developmental timing of expression or different effects in different tissues. It could also just reflect the variability inherent in these types of assays. Likewise, in a few cases, we detected an effect on STM but not LTM. While there are possible explanations for this, such as different temporal stages of memory engaging distinct neural circuits (Akalal et al., 2006; Blum et al., 2009; Wu et al., 2007) or different dynamics of neuromodulators (Aso et al., 2019, 2012; Aso and Rubin, 2016; Huetteroth et al., 2015; Yamagata et al., 2015), we are currently not interpreting these as bona fide phenotypes. We note too that in some cases, flies showed avoidance of the odor, rather than just loss of memory. Given the mechanism we uncovered, which suggests upregulation of Dop1R2 signaling in LL, we speculate that variations in the upregulation of Dop1R2 could account for this.

Our discovery that deviations in standard light exposure (ie. constant light or chronic jetlag light) result in impaired associative memory, which can be inherited by three generations of progeny raised in a standard light dark cycle, raises the question of whether similar phenomena and mechanisms can be extended to other behaviors and higher organisms. All molecular machinery implicated in this study have orthologs in mammals, and both piRNA and dopamine expression are affected by stressful environmental conditions. Indeed, chronic acute stress, such as dim light during dark phases, can result in dampened dopamine expression. Our findings highlight epigenetic modifications in response to environmental perturbations that impact a critical behavior (memory) across generations.

**Supplemental Information** Figures S1-S8

## ACKNOWLEDGEMENTS

We thank S. Killiany and E. Astacio for brain dissections, E. Astacio for ovary dissections, S. Killiany, E. Astacio, R. Xiong and P. Pivarshev for counting associative memory assays, K. Liuu for making Drosophila food, and the Sehgal lab for discussion. A.S. is an HHMI-Investigator. R.S.M. was supported by a HHMI-Damon Runyon Research Fellowship.

## AUTHOR CONTRIBUTIONS

Conceptualization, R.S.M., and A.S. Methodology, R.S.M., and A.S. Investigation, R.S.M., and A.S. Writing – Review and Editing, R.S.M., and A.S. Funding Acquisition, A.S.

---

**Supplement 1:**
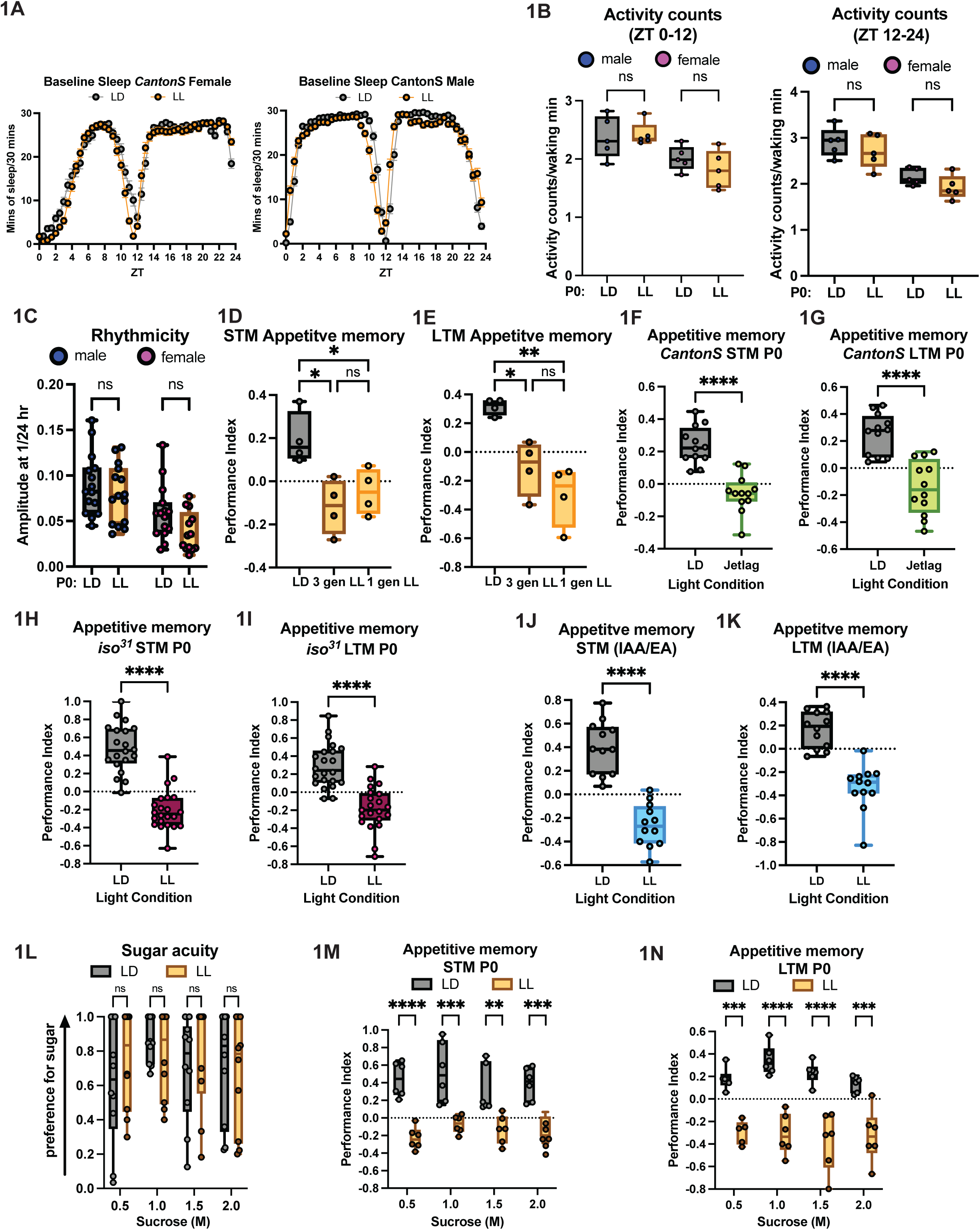
Effect of aberrant light on memory is generalizable to a chronic-jetlag paradigm as well as different odors and sugar concentrations (A) Mated female flies (left) or males (right) raised in LD or LL have comparable sleep profiles; n = 2 biological replicates, 16 flies per sex. (B) Mated female flies and males raised in LD or LL have similar activity during wake during the day (left, ZT 0-12) or night (right, ZT 12-24); n = 2 biological replicates, 16 flies per sex. (C) Mated females and males raised in LD and LL have similar behavioral rhythms in constant darkness; n = 2 biological replicates, 16 flies per sex. (D-E) One generation of LL is sufficient to impair short-term (D) and long-term (E) appetitive memory; n = 1 biological replicates, 4 replicate assays. (F-G) CantonS red eyed flies exposed to a jetlag light protocol (ie. 9 hr phase advance of light every 2 days) have impaired appetitive short-term (F) and long-term (G) memory performance; n = 3 biological replicates, 12 replicate assays. (H-I) *iso^31^* white eyed flies do not have standard aversive short-term (H) or long-term memory (I) in LL; n = 3 biological replicates, 20 replicate assays. (J-K) CantonS flies exposed to constant light cannot execute appetitive memory regardless of odor used (ie. IAA/EA); n = 2 biological replicates, 12 replicate assays. (L) Flies raised in LD or LL chemotaxis towards increasing concentrations of sugar; n = 2 biological replicate, 10 replicate assays. (M-N) Despite increasing concentrations of sugar flies exposed to constant light cannot perform appetitive short-term (M) or long-term (N) memory; n = 2 biological replicates, 6 replicate assays. (D-K) Unpaired t-test, (D-E, L-M) Two-way ANOVA, Tukey’s multiple comparison test, mean ± SEM. Each dot represents at least 50 flies. (B-C) One-way ANOVA, Tukey’s multiple comparison test, mean ± SEM. *p ≤ 0.05, **p ≤ 0.01, ***p ≤ 0.001, ****p ≤ 0.00001, ns, not significant.

**Supplement 2:**
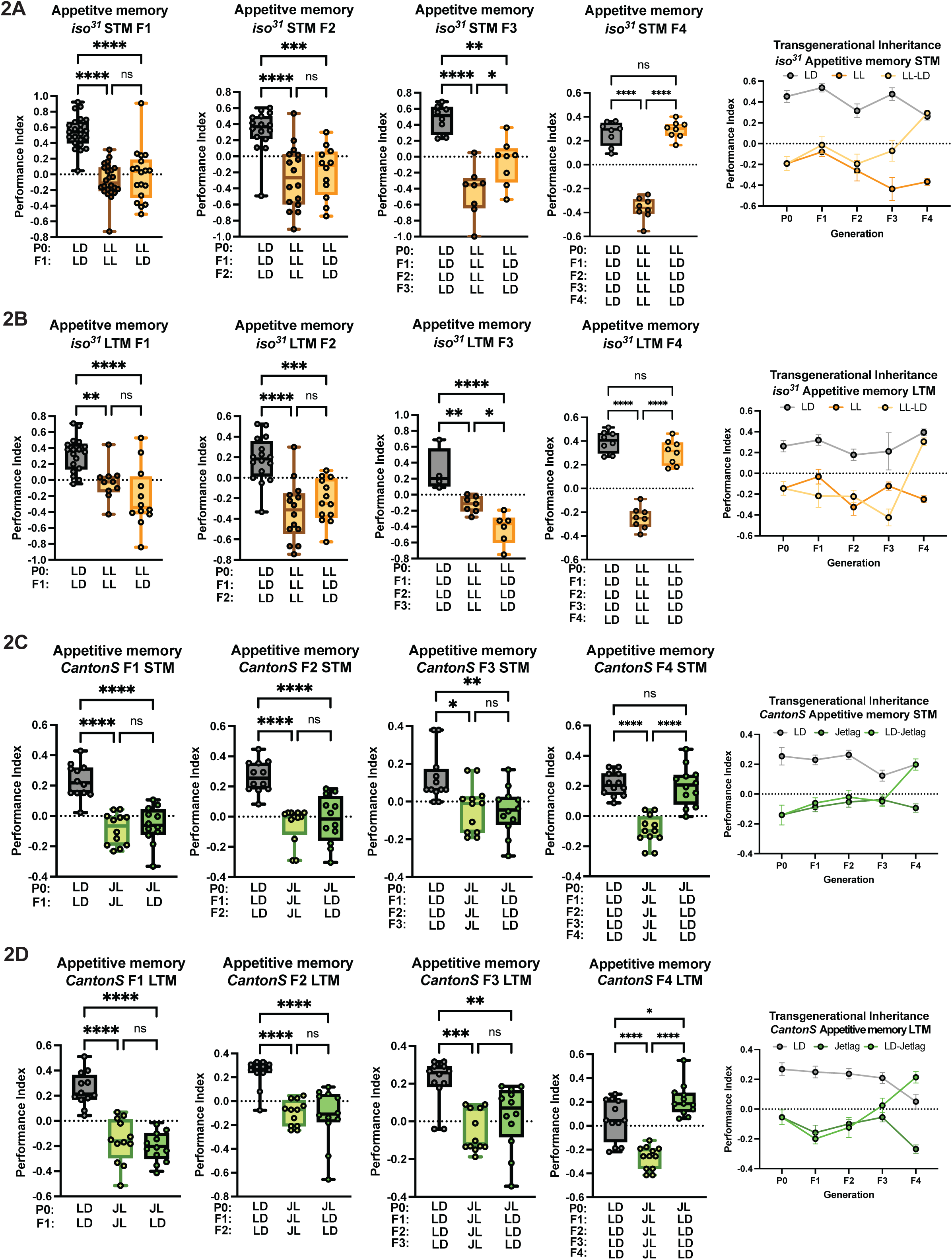
LL-impaired memory is transgenerationally inherited regardless of genotype or mode of light cycle disruption. (A-B) *iso^31^* white eyed flies’ progeny of LL-exposed parents transgenerationally inherit impaired short-term (A) and long-term (B) appetitive memory for three generations despite being raised in LD. Standard associative memory performance is restored in the fourth generation. n = 3 biological replicates, 18 replicate assays. (C-D) CantonS progeny of jet lag light (9 hr phase shift every 48 hours) exposed parents raised in LD have impaired appetitive short-term (C) and long-term (D) memory for three generations. n = 3 biological replicates, 12 replicate assays. Appetitive memory is restored in the fourth generation. One-way ANOVA, Tukey’s multiple comparison test, mean ± SEM. Each dot represents at least 50 flies. *p ≤ 0.05, **p ≤ 0.01, ***p ≤ 0.001, ****p ≤ 0.00001, ns, not significant.

**Supplement 3:**
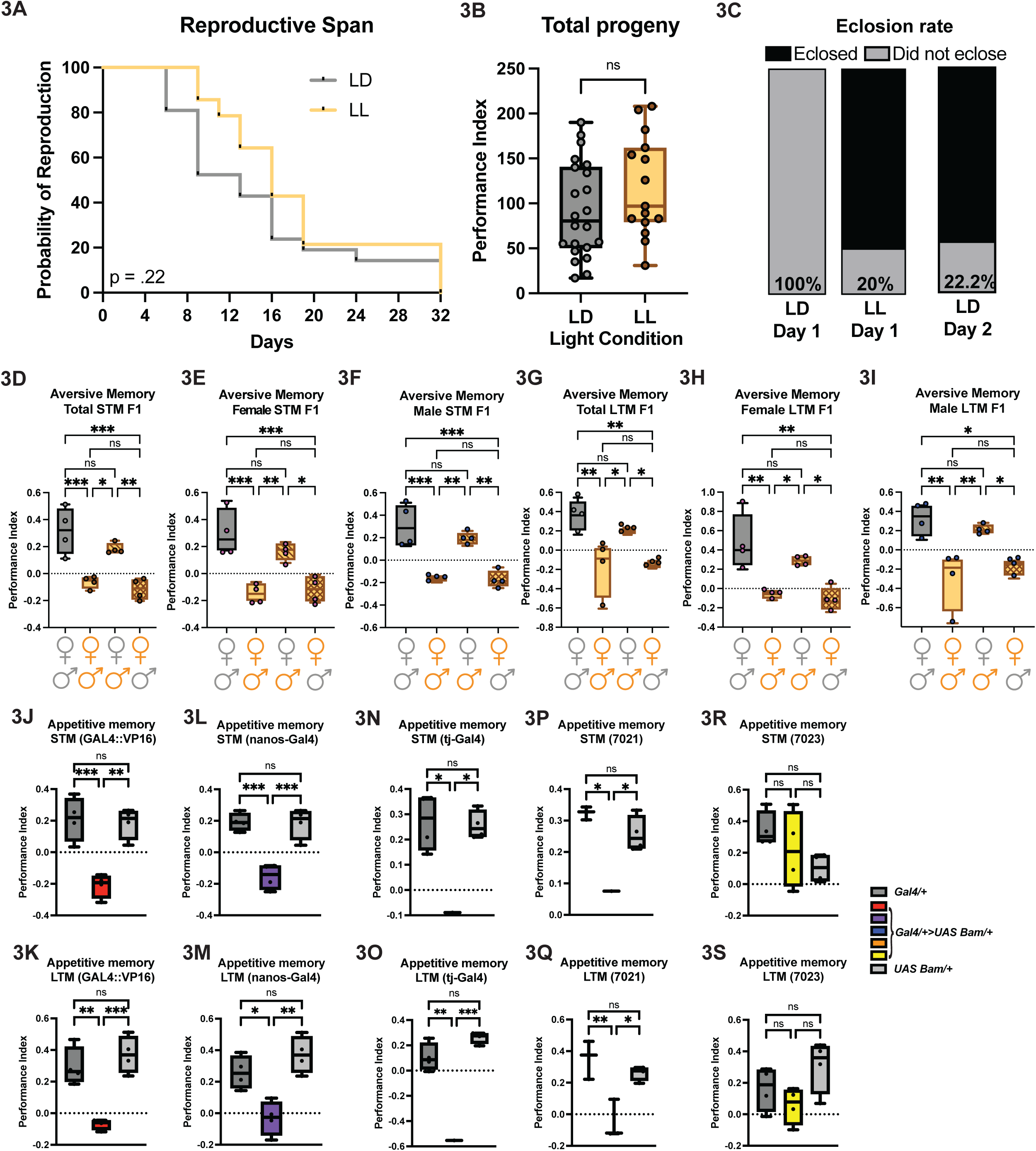
Irregular light exposure does not affect reproduction, but associative memory requires the germline. (A) No difference in reproductive span in flies exposed to LL compared to LD; n = 1 biological replicate, 24 flies. Log-Rank (Mantel-Cox test). (B) Mated females raised in LD and LL have the same number of total progeny eclose; n = 1 biological replicate, 10 flies. Student’s t-test. (C) Flies exposed to constant light eclose 24 hours earlier than LD flies (80% eclosed on Day 1 LL of eclosion vs 0% LD). n = 10 individual flies. (D-I) Progeny of LL-trained mated parents raised in LL (orange, LL males x orange, LD females) approach electric shock during short-term (D-F) and long-term (G-I) memory, while progeny of LD-trained mated parents raised in LD (gray, LD males x gray, LD females) avoid electric shock during short-term (D-F) and long-term (G-I) memory. Progeny of male parents exposed to LL inherit appropriate short-term (F-H) and long-term (G-I) memory, while progeny of mothers exposed to LL inherit impaired short-term (D-F) and long-term (G-I) memory; n = 1 biological replicates, 4 replicate assays. (J-S) *GAL4::VP16>UAS Bam* (J-K), *nanos-Gal4>UAS Bam* (L-M*), tj-Gal4>UAS Bam* (N-O), and *7021>UAS Bam* mated male and female flies (Red bar) impair appetitive STM (J, L, N, P) and LTM (K) in LD. (R-S*) 7023>UAS Bam* over expression does not affect STM (R) or LTM in LD. (S); n = 1 biological replicate, at least 2 replicate assays. One-way ANOVA, Tukey’s multiple comparison test, mean ± SEM. Each dot represents at least 50 flies. *p ≤ 0.05, **p ≤ 0.01, ***p ≤ 0.001, ns, not significant.

**Supplement 4:**
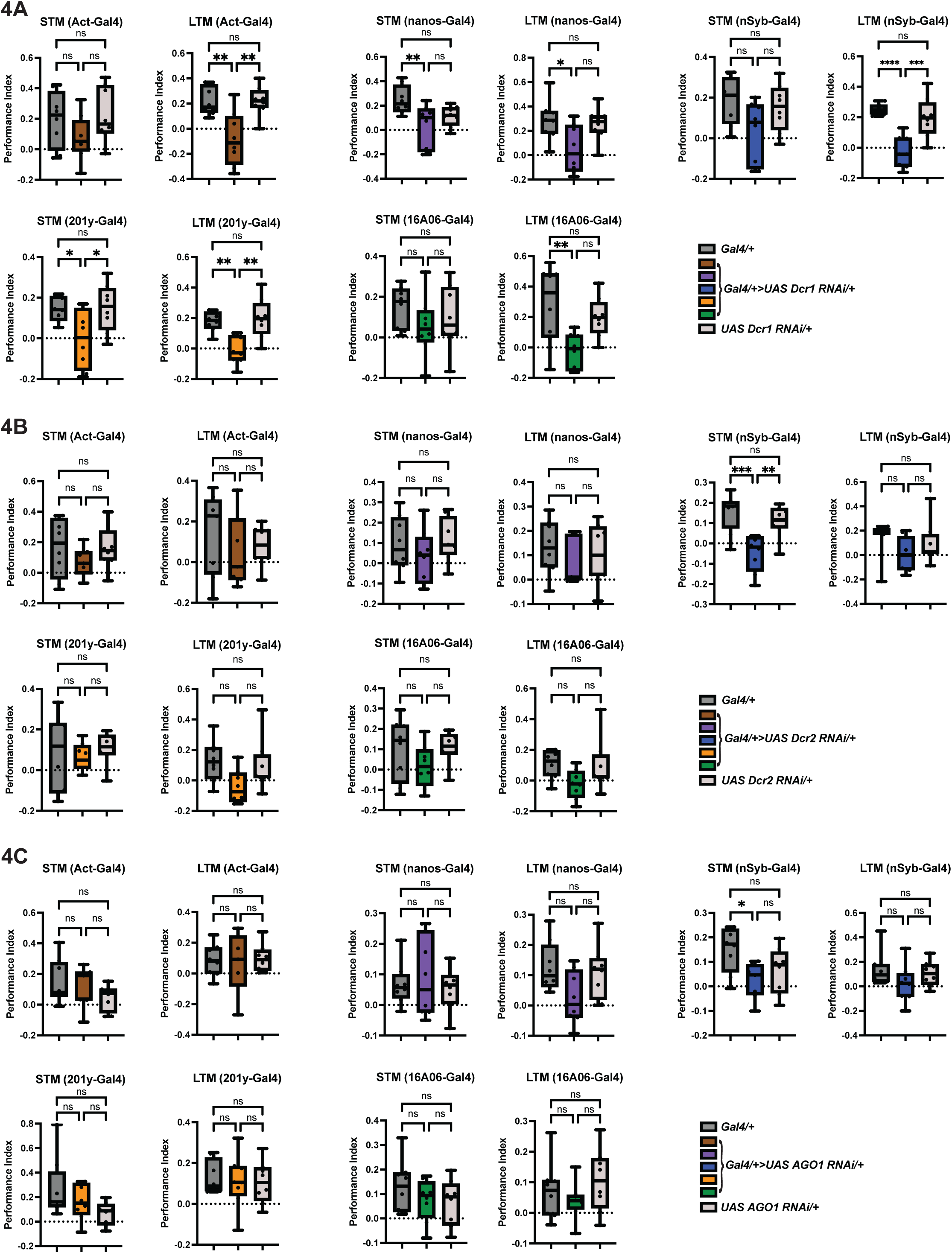
The miRNA and endo-siRNA pathways are not required for appetitive memory (A) Knockdown of Dcr1 in the mushroom body impairs STM and LTM, while knockdown of Dcr1 full body, pan neuronally, affects LTM in LD. (B) Dcr2 is only required pan neuronally for STM in LD. (C) AGO1 is not required in any tissue tested for STM and LTM in LD. n = 2 biological replicate, 8 replicate assays. Key: Brown = act-Gal4, purple = nanos-Gal4, blue = nSyb-Gal4, orange = 201y-Gal4, green = 16A06-Gal4. One-way ANOVA, Tukey’s multiple comparison test, mean ± SEM. Each dot represents at least 50 flies. *p ≤ 0.05, **p ≤ 0.01, ***p ≤ 0.001, ****p ≤ 0.0001 ns, not significant.

**Supplement 5:**
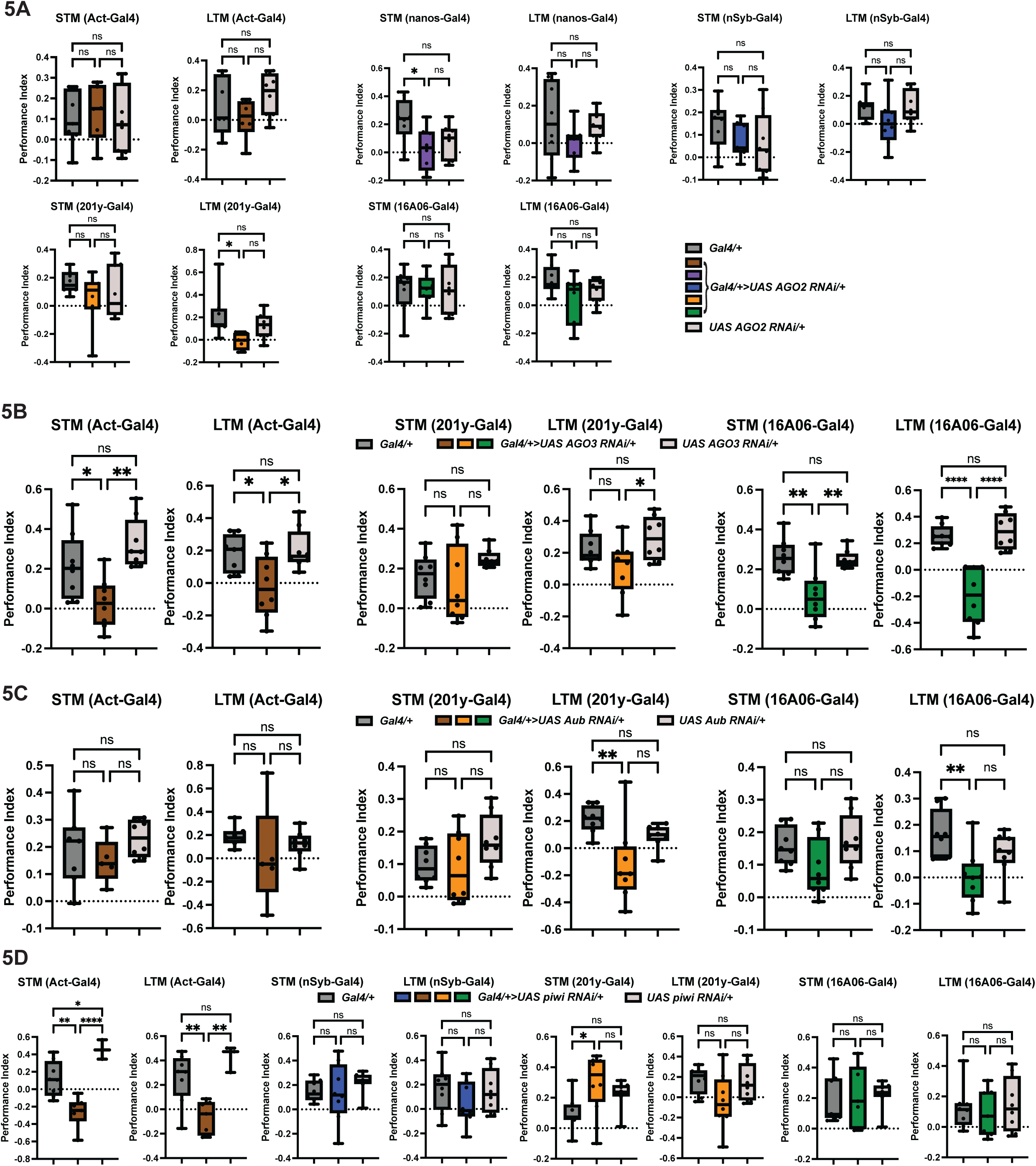
Piwi clade Argonautes are required for associative memory (A) AGO2 is not required in any tissue tested for appetitive STM or LTM in LD. (B) Knockdown of AGO3 full body or using the broad mushroom body driver impairs STM and LTM, while knock down in the mushroom body has no effect in LD. (C) Aub is not required full body, or in the mushroom body for STM or LTM in LD. (D) act-Gal4>UAS Piwi RNAi impairs appetitive STM and LTM in LD, while having no effect pan neuronally or in the mushroom body. n = 2 biological replicate, 8 replicate assays. Key: Brown = act-Gal4, purple = nanos-Gal4, blue = nSyb-Gal4, orange = 201y-Gal4, green = 16A06-Gal4. One-way ANOVA, Tukey’s multiple comparison test, mean ± SEM. Each dot represents at least 50 flies. *p ≤ 0.05, **p ≤ 0.01, ***p ≤ 0.001, ****p ≤ 0.0001 ns, not significant.

**Supplement 6:**
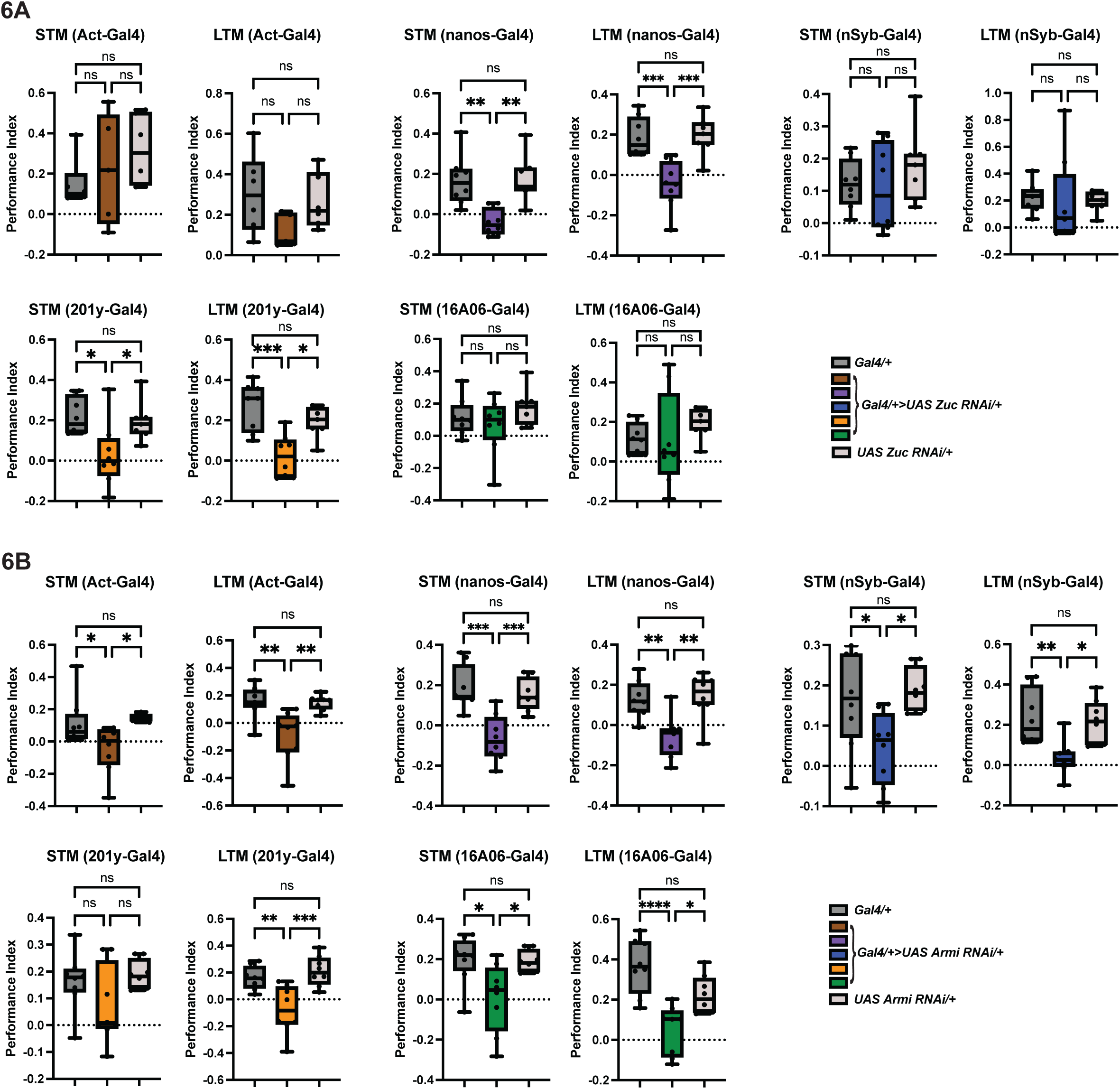
Primary piRNA biogenesis and the ping-pong cycle are required for associative memory (A) Knock down of Zuc in the germline or in the mushroom body impairs appetitive STM and LTM in LD. (B) Armi is broadly required for STM and LTM in LD: full body, germline, pan neuronally and in the mushroom body. n = 2 biological replicate, 8 replicate assays. Key: Brown = act-Gal4, purple = nanos-Gal4, blue = nSyb-Gal4, orange = 201y-Gal4, green = 16A06-Gal4. One-way ANOVA, Tukey’s multiple comparison test, mean ± SEM. Each dot represents at least 50 flies. *p ≤ 0.05, **p ≤ 0.01, ***p ≤ 0.001, ****p ≤ 0.0001 ns, not significant.

**Supplement 7:**
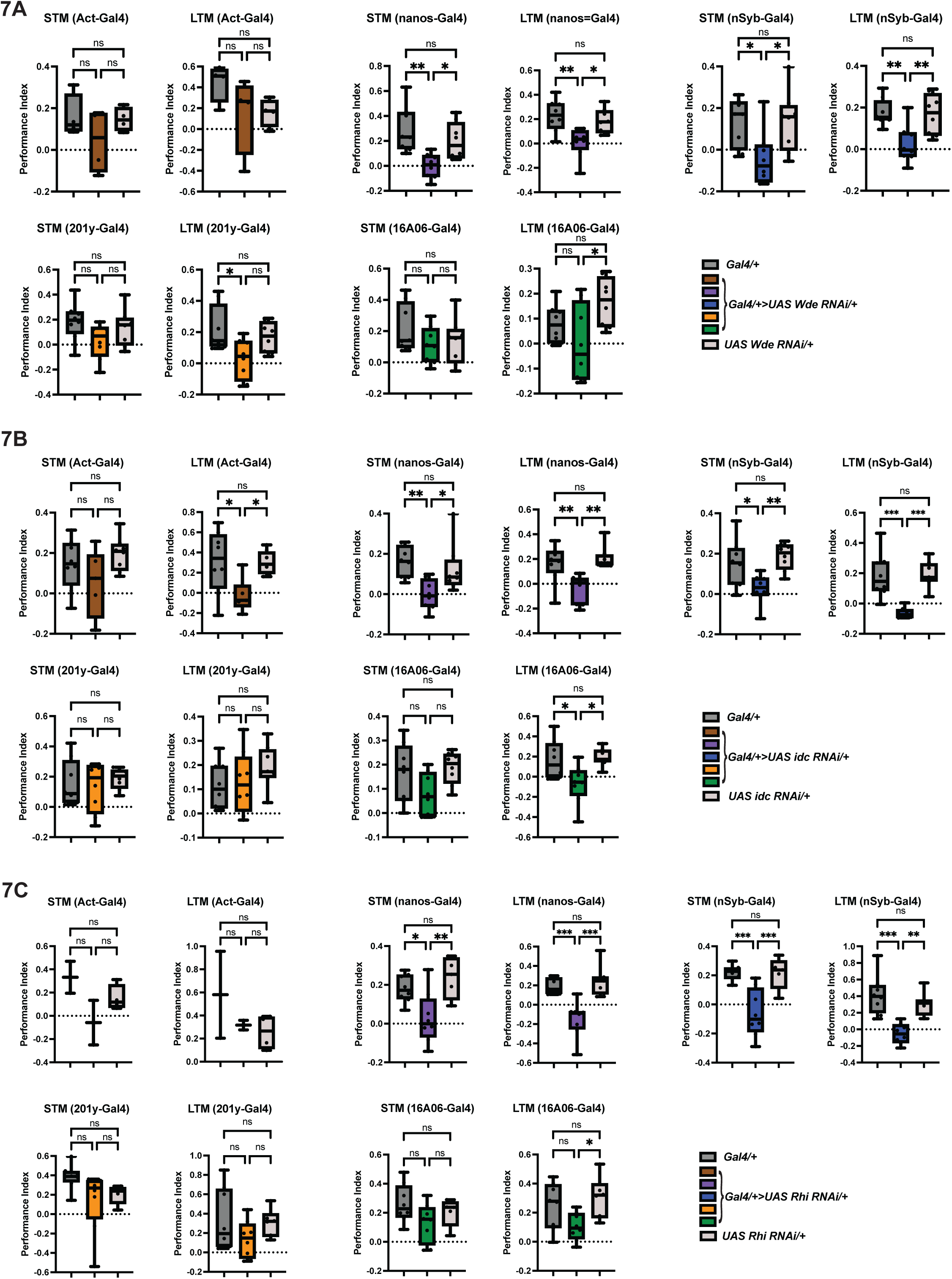
Deposition of H3K9me3 repressive chromatin marks is required for associative memory (A) *nanos-Gal4>UAS wde RNAi* (purple) and *nSyb-Gal4>UAS wde RNAi* (blue) invert appetitive STM and LTM in LD, while there is no effect when knocked down full body or in the mushroom body. (B) idc is required in the germline and neurons for standard appetitive STM and LTM, full body for LTM, and broadly in the mushroom body for STM. (C) Knockdown of Rhi in the germline or in neurons impairs appetitive STM and LTM. N = 2 biological replicate, 8 replicate assays. Key: Brown = act-Gal4, purple = nanos-Gal4, blue = nSyb-Gal4, orange = 201y-Gal4, green = 16A06-Gal4. One-way ANOVA, Tukey’s multiple comparison test, mean ± SEM. Each dot represents at least 50 flies. *p ≤ 0.05, **p ≤ 0.01, ***p ≤ 0.001, ****p ≤ 0.0001 ns, not significant.

**Supplement 8:**
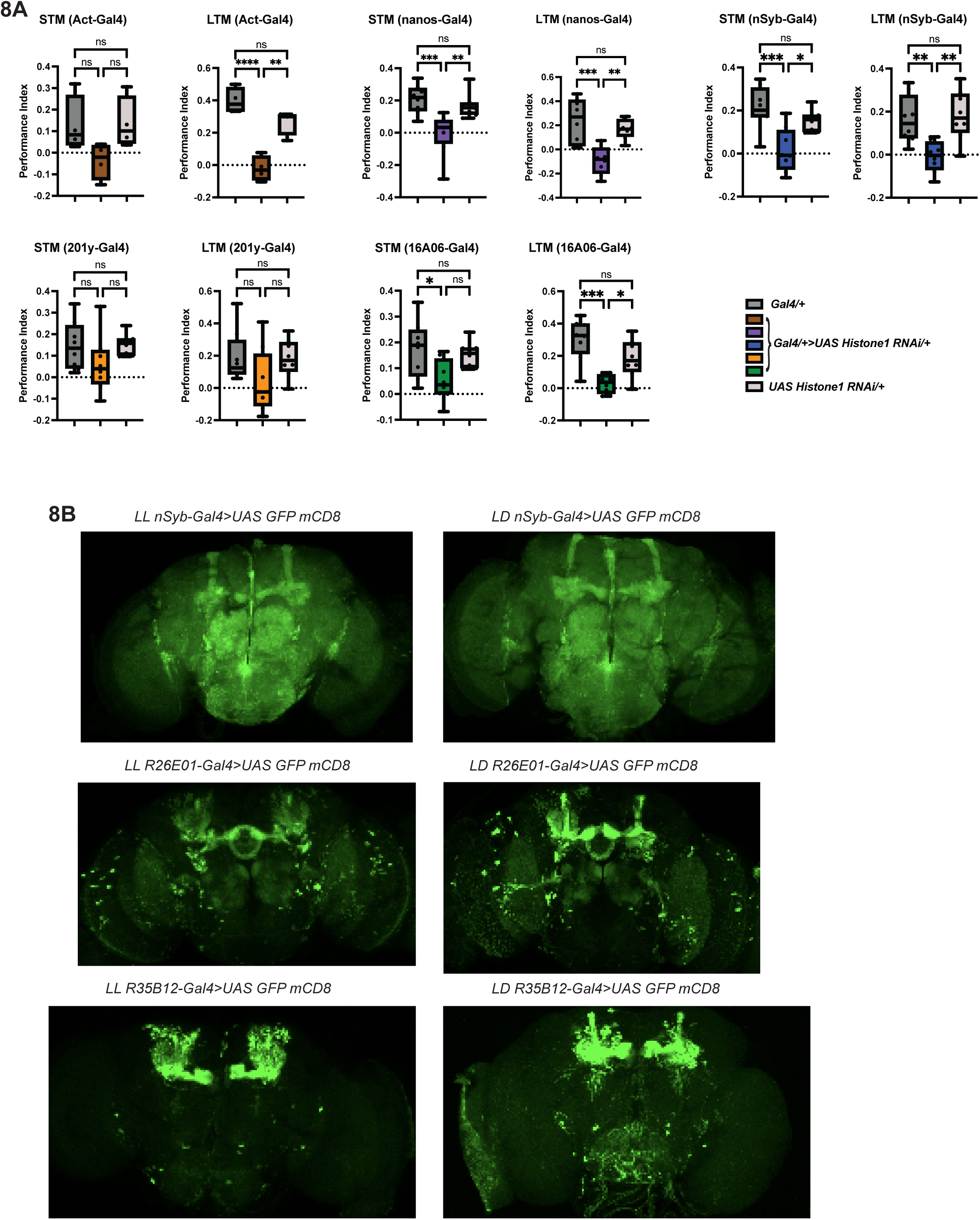
Effects of constant light are mediated by histone H1 but not through gross changes in brain architecture (A) Histone1 is required in the germline and neurons for standard appetitive STM and LTM, full body for LTM, and broadly in the mushroom body for STM in LD. n = 2 biological replicate, 8 replicate assays. Key: Brown = act-Gal4, purple = nanos-Gal4, blue = nSyb-Gal4, orange = 201y-Gal4, green = 16A06-Gal4. One-way ANOVA, Tukey’s multiple comparison test, mean ± SEM. Each dot represents at least 50 flies. *p ≤ 0.05, **p ≤ 0.01, ***p ≤ 0.001, ****p ≤ 0.0001 ns, not significant. (B) Constant light does not change overall brain architecture (top, *nSyb-Gal4>UAS GFP mCD8*), or mushroom body architecture (middle, *R26E01-Gal4>UAS GFP* mCD8, bottom, *R35B12>UAS GFP mCD8*). n = 2 biological replicates, 10 fly brains.

## EXPERIMENTAL MODEL AND STUDY PARTICIPANT DETAILS

### Drosophila melanogaster

RNAi and Gal4 lines were ordered from Bloomington Drosophila Stock Center in Indiana, or Vienna Drosophila Resource Center in Austria (information about individuals lines can be found in the key resource table). *piwi^1^ and piwi^2^* were generously gifted by Haifin Lin for memory assays (Lin and Spradling, 1997). *per^0^* (Benzer lab) and *tim^01^* (Young lab), cry^b^ (Hall lab) have been in our lab for many years. Iso^31^, CantonS Red eye, yw, w^1118^, and nSyb-Gal4 were acquired from lab stocks.

*Piwi^1^*; UAS Tango Dop1R1 flies were made by mating *piwi^1^* x Bloomington Stock Center line 68234.

To knock down maternal piRNAs female *piwi^1^* heterozygotes *and piwi^2^* homozygotes were mated to control males.

Adult flies were raised in 12:12 LD light:dark cycle incubators, 12:12 LL light:light cycle incubators, or in 12:12 LD light:dark cycles with the phase of light advanced by 9 hours every 48 hours. All cultures were at ∼25°C and ∼65% humidity.

Four- to seven-day-old flies were used for all behavior experiments and were transferred to fresh food vials 48 hours before behavioral tests and dissections. Mated flies were used for all studies, except for Figure 2D and 2E, all of which were collected and verified to be virgins. All flies used in these experiments were aged to day 4-7 of adulthood prior to experimentation. The fly population was randomized but was kept age matched in each trial. For food deprivation prior to behavioral assays, flies were kept in vials with 2% agar to prevent desiccation.

Flies were maintained on standard cornmeal-molasses diet, including 64.7g/L cornmeal, 27.1 g/L dry yeast, 8g/L agar, 61.6 mL/L molasses, 10.2 mL/L 20% tegosept, and 2.5 mL/L propionic acid.

## METHODS

### Dissections

At eclosion virgin females were collected and mated until day 4-7 of adulthood.

Brain dissections: Flies were appetitive memory trained or mock trained (odor presentation only) immediately before dissection. Flies were anesthetized on ice and dissecting dishes were cooled on ice prior to dissections. Fly brains were dissected in cold 1x PBS and fixed in 4% paraformaldehyde (PFA) for 30-mins rocking at room temperature. Brains were washed 3 x 15 minutes with PBS with 0.4% Triton-X100 (PBST) rocking at room temperature and mounted with Vectashield. Brains were imaged using a Leica Stellaris 8 TauSTED confocal microscope. FIJI software was used for analysis.

Ovary dissections: Flies were anesthetized on ice and dissecting dishes were cooled on ice prior to dissections. Fly ovaries were dissected in cold Schneider’s Drosophila Media. Immediately after dissection individual ovarioles were separated using a 22G needle and then fixed in 4% paraformaldehyde (PFA) for 30-mins rocking at room temperature. Ovaries were washed 3 x 15 minutes in Schneider’s rocking at room temperature. Following the final wash ovaries were quick washed in 1x PBS. Ovaries were mounted in Vectashield. Ovaries were imaged using a Leica Stellaris 8 TauSTED confocal microscope. FIJI software was used for analysis.

In experiments in which F1 and subsequent generations were used: Some of the pooled animals were subjected to a behavior assay, while the remaining were moved into new bottles for 36-hour egg-lays before disposing of parents. Half of the P0 LL egg-lay was moved to 12:12 LD for the rest of the experiment.

### Imaging and Fluorescence Quantitation

Brains: For all representative images, for visualization purposes, brightness and contrast were adjusted identically across groups. Z-stacks of brains were imaged every 2 μm at 20x magnification; maximum intensity projections of brains were built with Fiji. To quantify total fluorescence ROIs were drawn around the brains. To quantify number of puncta maximum intensity projections of brains were built with Fiji. Images were thresholded using IJ-isodata with smoothing and were adjusted to visualize all puncta and remove background noise. Images were then made binary and converted to a mask, with watershed. Puncta were quantified using a pixel size range of 8 to infinity.

Ovaries: Z-stacks of ovaries were imaged every 3 μm at 10x magnification; maximum intensity projections of brains were built with Fiji. To quantify nuclear fluorescence, ROIs were drawn around individual egg chambers and total fluorescence per egg chamber was divided by the number of nuclei.

### Behavior

Appetitive short-term conditioning was performed as described previously (Colomb et al., 2009; Krashes et al., 2009). In brief, a 4- to 7-day-old mixed sex population of ∼100 flies was starved for 18 hours in vials containing 2% agar and then trained at 25°C and 70% relative humidity in the dark with a red light to associate 1.5 M sucrose (soaked on 1 in x 1 in squares of Whatman paper dried with a blow dryer) (unconditioned stimulus, CS) with odor A (conditioned stimulus, US), presented in an air stream, for 2 minutes followed by a 30-second stream of clean air. Then flies were presented with a water-soaked Whatman paper (blank) plus odor B (CS) for 2 minutes, followed by another 30-second stream of clean air. In reciprocal experiments, odor B and odor A were presented with sucrose and the blank, respectively.

Aversive short-term conditioning was performed as described previously (Malik and Hodge, 2014; Tully and Quinn, 1985). In brief, a 4- to 7-day-old mixed sex population of ∼100 flies were starved for 18 hours in vials containing 2% agar and then trained at 25°C and 70% relative humidity in the dark with a red light to avoid odor A that is paired with an electric shock (US, oscillator settings: frequency 2 x .1 s, duration 20 x 10, delay 20 x 10), followed by a 30-second stream of clean air. Next, flies were presented with odor B with no electric shock (CS), followed by another 30-second stream of clean air. In reciprocal experiments, odor B and odor A were presented with electric shock and the blank, respectively.

During both short-term and long-term memory training and testing all flies were kept in the room in which memory experiments are being conducted (i.e. darkness with a red light) for the duration of the experiment (at most 6 hours). Immediately following short-term memory, flies were refed normal food for at least three hours. During this time flies were anesthetized and counted on CO_2_ pads. Flies were kept on the bench during this time. Following at least three hours of refeeding flies were starved 16-18 hours overnight in vials on 2% agar to perform long-term memory. Long-term memory was tested by presenting flies in a T-maze with odor A and odor B for 2 minutes

The odors used in these experiments were: 4-methylcyclohexanol paired with 3-octanol, or ethyl acetate paired with isoamyl acetate. All odors were diluted in paraffin oil: 4-methycyclohexanol was diluted 1:200, 3-octanol was diluted 1:80, ethyl acetate was diluted to 1:200, and isoamyl acetate was diluted to 1:75. Odors were presented in 3-mm diameter cups in the airstream. The performance index was calculated as the number of flies selecting US odor minus the number of flies selecting the CS+ odor divided by the total number of flies. Each performance index is the average of the performance indices from reciprocal experiments with two odors swapped to minimize non-associative effects.

In experiments in which F1 and subsequent generations were used: Some of the pooled animals were subjected to a behavior assay, while the remaining were moved into new bottles for 36-hour egg-lays before disposing of parents. Half of the P0 LL egg-lay was moved to 12:12 LD for the rest of the experiment.

### Sugar Acuity

4-7 day old mixed sex populations of ∼100 flies were starved for 18 hours in vials on 2% agar and then trained at 25°C and 70% relative humidity in the dark with a red light and given the choice between increasing concentrations of sucrose (0.5 M, 1.0 M, 1.5 M, 2.0 M) or water. Flies were anesthetized and counted on CO_2_ pads. Performance indices were calculated as described in behavior.

### Reproductive span and progeny count

Mated reproductive spans were performed as previously described (Von Philipsborn et al., 2023). In brief virgin females were collected in groups and aged to day 2 of adulthood. At day 2 individual females were moved into individual food vials and mated with one male for 24 hours. Males were removed after mating. Mated females were moved into new vials daily until the end of the reproductive span. 48 hours after removal, vials were screened for progeny to confirm egg production. Reproductive cessation was defined as the last day of progeny production preceding two full days without progeny. For a subset of mated females, progeny vials were maintained until eclosion. Vials were counted every day to determine number of eclosed progeny.

### Virgin behavior assay

Male and female virgins were collected, raised in 12:12 LD or LL, and aged to day 4-7 of adulthood before performing short- and long-term appetitive memory as described above. Following behavior confirmation, the following matings were set up: 1. LD males x LD females, 2. LL males x LL females, 3. LD males x LL females, and 4. LL males x LD females. Group 1, 3, and 4 were maintained in LD, while group 2 was maintained in LL. Groups were left to mate for 36 hours prior to removal of parents. Subsets of F1 progeny were aged to day 4-7 of adulthood before performing appetitive and aversive short- and long-term appetitive memory as described above.

### Sleep Assay

For all sleep experiments single beam *Drosophila* Activity Monitoring System (DAM2, Trikinetics) was used as previously described (Hendricks et al., 2000). Briefly, 4-7 day old, mated females and males that had been raised in either 12 hr: 12 hr light-dark or 12 hr: 12 hr light-light were loaded into 2×60 mm glass locomotor tubes, using DAM2 *Drosophila* activity monitors from Trikinetics. Locomotor tubes were loaded with 2% agar with 5% sucrose as fly food on one side and yarn on the other to plug the tube. Montiors were placed in incubators kept at 25°C in a 12 hr:12 hr light-dark cycle and 50% humidity. In this assay, activity counts correspond to breaks in one infrared beam that bisect the tube. Sleep is defined as an activity count of zero for a minimum of 5 consecutive minutes. Five consecutive days were used for sleep and activity during time awake analyzed by Cynthesis (Sehgal Lab Matlab program).

### Rest:Activity Rhythm Assay

Locomotor activity assays were performed with the *Drosophila* Activity Monitoring System (Trikinetics) as described previously reported (Williams et al., 2001). Flies were raised and entrained to a 12 hr:12 hr light-dark cycle or 12 hr: 12 hr light-light cycle until day 4-7 of adulthood. 4-7 day old. Individual mated male and female flies were loaded into 2×60 mm glass tubes containing 5% sucrose and 2% agar as fly food on one side and yarn on the other side to plug the tube. Loaded tubes were transferred to constant darkness for 7 days at 25°C. Rhythm strength was determined by FFT analysis. A fly was considered rhythmic if it met two criteria: (1) displayed a rhythm with 95% confidence using χ^2^ periodogram analysis and (2) a corresponding FFT value above 0.01. Some flies exhibited bimodal periodicity, we took the larger FFT value for the determined period of the first harmonic of that period. Rhythm strength was categorized as weak (0.01-0.03), moderate (0.03-0.05), or strong (≥0.05). Locomotor activity was analyzed with ClockLab software (Actimetrics, Wilmette IL).

## QUANTIFICATION AND STATISTICAL ANALYSIS

The statistical details of experiments can be found in the figure legends. Reproductive spans were assessed using Kaplan-Meier log rank tests. For comparison of performance indices, FFTs, or waking activity between genotypes one-way ANOVA with Tukey’s multiple comparison test were used. For the comparison of performance indices between two conditions (i.e. 12:12 LD versus LL), of brain fluorescence, puncta count, or progeny eclosion rates, unpaired t-tests were performed. Experiments were repeated on separate days with separate populations, to confirm that results were reproducible. All images within groups were taken on the same day to exclude any differences in microscope operation. Flies of both sexes were used in sleep and behavioral rhythm experiments but were analyzed separated due to the known differences in sleep behavior between male and female flies using a Sehgal Lab analysis pipeline Prism 8 software was used for all statistical analyses. Additional, statistical details of experiments, including sample size (with n representing the number of behavior assays performed for appetitive and aversive memory, number of individual brains or ovaries images, and number of flies used in these assays) can be found in the figure legends.

### Key resources table

| REAGENT or RESOURCE | SOURCE | IDENTIFIER |
| --- | --- | --- |
| Agar, Drosophila Type | Fisher | Cat# NC1429200 |
| Agar | EMD Millipore Corp | Cat# 12177 |
| Molasses | Genessee Scientific | Cat# 62-118 |
| Corn Meal | LabScientific | Cat# FLY-8010-20 |
| Drosophila Dry Active Yeast | LabScientific | Cat# FLY-8040-10 |
| Schneiders Drosophila Medium | Life Technologies | Cat# 21720024 |
| D-Sucrose | Fisher Scientific | Cat# BP220-212 |
| Paraffin Oil | Sigma Aldrich | Cat# 8012-95-1 |
| 1x Phosphate Buffered Saline | Fisher Scientific | Cat# 14190235 |
| Triton-x100 | Sigma Aldrich | Cat# 9036-19-5 |
| Whatman Paper (3 MM) | Cytiva | Cat# 3030-662 |
| 4-Methylcyclohexanol | Sigma Aldrich | Cat# 589-91-3 |
| 3-Octanol | Sigma Aldrich | Cat# 589-98-0 |
| Ethyl Acetate | Fisher Chemical | Cat# 141-78-6 |
| Isoamyl Acetate | ThermoScientific | Cat# 123-92-2 |
| Paraformaldehyde (16%) | Electron Microscopy Sciences | Cat# 15710-S |
| VectaShield Mounting Medium | Vector Labs | Cat# H-1000 |
| Tegosept | Sigma Aldrich | Cat# 3647 |
| Experimental models: Organisms/strains |  |  |
| Iso31 | Lab generated strain |  |
| CantonS Red eye | Lab generated strain |  |
| per<sup>0</sup> | Gift of Seymour Bnezer | Benzer, 1971 |
| tim<sup>01</sup> | Gift of Michael Young | Sehgal et al. 1994 |
| glass<sup>60j</sup> | Bloomington Stock Center | Cat# 509 |
| cry<sup>b</sup> | Gift of Jeff Hall | Stanewsky et al. 1998 |
| UAS Bam | Bloomington Stock Center | Cat# 33631 |
| GAL4::VP16 | Bloomington Stock Center | Cat# 4937 |
| tj-Gal4 | Vienna Drosophila Resource Center | Cat# 104034 |
| P{w[+mW.hs]=GawB}112A | Bloomington Stock Center | Cat# 7021 |
| nanos-Gal4 | Bloomington Stock Center | Cat# 25721 |
| y[1] w[*]; P{w[+mW.hs]=GawB}109-30/CyO | Bloomington Stock Center | Cat# 7023 |
| UAS AGO3 RNAi | Bloomington Stock Center | Cat# 34815 |
| UAS Aub RNAi | Bloomington Stock Center | Cat# 39026 |
| UAS Piwi RNAi | Bloomington Stock Center | Cat# 37483 |
| Act-Gal4 | Bloomington Stock Center | Cat# 25374 |
| nSyb-Gal4, outcrossed 5x to iso31 | Lab generated allele |  |
| 201y-Gal4 | Lab generated allele/Available at Bloomington Stock Center | Cat# 4440, Crocker et al. 2010 |
| 16A06-Gal4 | Bloomington Stock Center | Cat# 48709 |
| UAS Dcr2 RNAi | Bloomington Stock Center | Cat# 27486 |
| UAS Zuc RNAi | Bloomington Stock Center | Cat# 36742 |
| UAS AGO1 RNAi | Bloomington Stock Center | Cat# 33727 |
| UAS armi RNAi | Bloomington Stock Center | Cat# 34789 |
| piw<sup>i1</sup> | Gift of Haifin Lin | Lin and Spradling 1997 |
| piw<sup>i2</sup> | Gift of Haifin Lin | Lin and Spradling 1997 |
| UAS Egg RNAi | Bloomington Stock Center | Cat# 36797 |
| UAS HP1 RNAi | Bloomington Stock Center | Cat# 36792 |
| UAS wde RNAi | Bloomington Stock Center | Cat# 33339 |
| UAS idc RNAi | Bloomington Stock Center | Cat# 64550 |
| UAS Rhi RNAi | Bloomington Stock Center | Cat# 34071 |
| EGFP::Piwi | Vienna Drosophila Resource Center | Cat# 313319 |
| R26E01-Gal4 | Bloomington Stock Center | Cat# 60510 |
| R35B12-Gal4 | Bloomington Stock Center | Cat# 49822 |
| UAS GFP mCD8 | Bloomington Stock Center | Cat# 5137 |
| UAS Histone1 RNAi | Bloomington Stock Center | Cat# 35205 |
| UAS Dop1R2 RNAi | Bloomington Stock Center | Cat# 51423 |
| UAS Dop1R1 Rango | Bloomington Stock Center | Cat# 68234 |
| UAS AGO2 RNAi | Bloomington Stock Center | Cat# 34799 |
| UAS Dcr1 RNAi | Bloomington Stock Center | Cat# 42901 |
| lexAop(-FRT)tdTomato.nls | Bloomington Stock Center | Cat# 66690 |
| Yw | Lab generated allele |  |
| W1118 | Lab generated allele |  |
| Software and algorithms |  |  |
| GraphPad Prism | GraphPad | Download at https://graphpad.com |
| Fiji | Fiji | Download at https://imagej.net/software/fiji |
| ClockLab | Actimetrics | Download at https://actimetrics.com/products/clocklab/clocklab-analyses/ |
| MatLab | Mathworks | Download at https://mathworks.com |
| DAM File Scan | Trikinetics | Download at https://trikinetix.com |
| Other |  |  |
| Oscilloscope | GW Instek | Cat# GOS-6112 |
| Single-Beam Sleep Monitors | Trikinetics | DAMH2 |

## REFERENCES

Adan, A., Archer, S.N., Hidalgo, M.P., Di Milia, L., Natale, V., Randler, C., 2012. Circadian Typology: A Comprehensive Review. Chronobiol. Int. 29, 1153–1175. 10.3109/07420528.2012.719971

Agrawal, P., Houl, J.H., Gunawardhana, K.L., Liu, T., Zhou, J., Zoran, M.J., Hardin, P.E., 2017. Drosophila CRY Entrains Clocks in Body Tissues to Light and Maintains Passive Membrane Properties in a Non-clock Body Tissue Independent of Light. Curr. Biol. 27, 2431–2441.e3. 10.1016/j.cub.2017.06.064

Akalal, D.-B.G., Wilson, C.F., Zong, L., Tanaka, N.K., Ito, K., Davis, R.L., 2006. Roles for *Drosophila* mushroom body neurons in olfactory learning and memory. Learn. Mem. 13, 659–668. 10.1101/lm.221206

Arai, J.A., Li, S., Hartley, D.M., Feig, L.A., 2009. Transgenerational Rescue of a Genetic Defect in Long-Term Potentiation and Memory Formation by Juvenile Enrichment. J. Neurosci. 29, 1496–1502. 10.1523/JNEUROSCI.5057-08.2009

Ashe, A., Sapetschnig, A., Weick, E.-M., Mitchell, J., Bagijn, M.P., Cording, A.C., Doebley, A.-L., Goldstein, L.D., Lehrbach, N.J., Le Pen, J., Pintacuda, G., Sakaguchi, A., Sarkies, P., Ahmed, S., Miska, E.A., 2012. piRNAs Can Trigger a Multigenerational Epigenetic Memory in the Germline of C. elegans. Cell 150, 88–99. 10.1016/j.cell.2012.06.018

Aso, Y., Herb, A., Ogueta, M., Siwanowicz, I., Templier, T., Friedrich, A.B., Ito, K., Scholz, H., Tanimoto, H., 2012. Three Dopamine Pathways Induce Aversive Odor Memories with Different Stability. PLoS Genet. 8, e1002768. 10.1371/journal.pgen.1002768

Aso, Y., Ray, R.P., Long, X., Bushey, D., Cichewicz, K., Ngo, T.-T., Sharp, B., Christoforou, C., Hu, A., Lemire, A.L., Tillberg, P., Hirsh, J., Litwin-Kumar, A., Rubin, G.M., 2019. Nitric oxide acts as a cotransmitter in a subset of dopaminergic neurons to diversify memory dynamics. eLife 8, e49257. 10.7554/eLife.49257

Aso, Y., Rubin, G.M., 2016. Dopaminergic neurons write and update memories with cell-type-specific rules. eLife 5, e16135. 10.7554/eLife.16135

Bagijn, M.P., Goldstein, L.D., Sapetschnig, A., Weick, E.-M., Bouasker, S., Lehrbach, N.J., Simard, M.J., Miska, E.A., 2012. Function, Targets, and Evolution of *Caenorhabditis elegans* piRNAs. Science 337, 574–578. 10.1126/science.1220952

Bannister, A.J., Zegerman, P., Partridge, J.F., Miska, E.A., Thomas, J.O., Allshire, R.C., Kouzarides, T., 2001. Selective recognition of methylated lysine 9 on histone H3 by the HP1 chromo domain. Nature 410, 120–124. 10.1038/35065138

Barrio-Alonso, E., Lituma, P.J., Notaras, M.J., Albero, R., Bouchekioua, Y., Wayland, N., Stankovic, I.N., Jain, T., Gao, S., Calderon, D.P., Castillo, P.E., Colak, D., 2023. Circadian protein TIMELESS regulates synaptic function and memory by modulating cAMP signaling. Cell Rep. 42, 112375. 10.1016/j.celrep.2023.112375

Benzer, S., 1971. From the Gene to Behavior. JAMA J. Am. Med. Assoc. 218, 1015. 10.1001/jama.1971.03190200047010

Berry, J.A., Phan, A., Davis, R.L., 2018. Dopamine Neurons Mediate Learning and Forgetting through Bidirectional Modulation of a Memory Trace. Cell Rep. 25, 651–662.e5. 10.1016/j.celrep.2018.09.051

Blum, A.L., Li, W., Cressy, M., Dubnau, J., 2009. Short- and long-term memory in Drosophila require cAMP signaling in distinct neuron types. Curr. Biol. CB 19, 1341–1350. 10.1016/j.cub.2009.07.016

Bozler, J., Kacsoh, B.Z., Bosco, G., 2019. Transgenerational inheritance of ethanol preference is caused by maternal NPF repression. eLife 8, e45391. 10.7554/eLife.45391

Brennecke, J., Malone, C.D., Aravin, A.A., Sachidanandam, R., Stark, A., Hannon, G.J., 2008. An Epigenetic Role for Maternally Inherited piRNAs in Transposon Silencing. Science 322, 1387–1392. 10.1126/science.1165171

Brower-Toland, B., Findley, S.D., Jiang, L., Liu, L., Yin, H., Dus, M., Zhou, P., Elgin, S.C.R., Lin, H., 2007. *Drosophila* PIWI associates with chromatin and interacts directly with HP1a. Genes Dev. 21, 2300–2311. 10.1101/gad.1564307

Casier, K., Delmarre, V., Gueguen, N., Hermant, C., Viodé, E., Vaury, C., Ronsseray, S., Brasset, E., Teysset, L., Boivin, A., 2019. Environmentally-induced epigenetic conversion of a piRNA cluster. eLife 8, e39842. 10.7554/eLife.39842

Castillo Díaz, F., Caffino, L., Fumagalli, F., 2021. Bidirectional role of dopamine in learning and memory-active forgetting. Neurosci. Biobehav. Rev. 131, 953–963. 10.1016/j.neubiorev.2021.10.011

Chellappa, S.L., Steiner, R., Blattner, P., Oelhafen, P., Götz, T., Cajochen, C., 2011. Non-Visual Effects of Light on Melatonin, Alertness and Cognitive Performance: Can Blue-Enriched Light Keep Us Alert? PLoS ONE 6, e16429. 10.1371/journal.pone.0016429

Chen, Q., Yan, M., Cao, Z., Li, X., Zhang, Yunfang, Shi, J., Feng, G., Peng, H., Zhang, X., Zhang, Ying, Qian, J., Duan, E., Zhai, Q., Zhou, Q., 2016a. Sperm tsRNAs contribute to intergenerational inheritance of an acquired metabolic disorder. Science 351, 397–400. 10.1126/science.aad7977

Chen, Q., Yan, M., Cao, Z., Li, X., Zhang, Yunfang, Shi, J., Feng, G., Peng, H., Zhang, X., Zhang, Ying, Qian, J., Duan, E., Zhai, Q., Zhou, Q., 2016b. Sperm tsRNAs contribute to intergenerational inheritance of an acquired metabolic disorder. Science 351, 397–400. 10.1126/science.aad7977

Crocker, A., Shahidullah, M., Levitan, I.B., Sehgal, A., 2010. Identification of a neural circuit that underlies the effects of octopamine on sleep:wake behavior. Neuron 65, 670–681. 10.1016/j.neuron.2010.01.032

De Belle, J.S., Heisenberg, M., 1994. Associative Odor Learning in *Drosophila* Abolished by Chemical Ablation of Mushroom Bodies. Science 263, 692–695. 10.1126/science.8303280

Dias, B.G., Ressler, K.J., 2014. Parental olfactory experience influences behavior and neural structure in subsequent generations. Nat. Neurosci. 17, 89–96. 10.1038/nn.3594

Dietz, D.M., LaPlant, Q., Watts, E.L., Hodes, G.E., Russo, S.J., Feng, J., Oosting, R.S., Vialou, V., Nestler, E.J., 2011. Paternal Transmission of Stress-Induced Pathologies. Biol. Psychiatry 70, 408–414. 10.1016/j.biopsych.2011.05.005

Dubnau, J., Grady, L., Kitamoto, T., Tully, T., 2001. Disruption of neurotransmission in Drosophila mushroom body blocks retrieval but not acquisition of memory. Nature 411, 476–480. 10.1038/35078077

Emery, P., Stanewsky, R., Hall, J.C., Rosbash, M., 2000. A unique circadian-rhythm photoreceptor. Nature 404, 456–457. 10.1038/35006558

Erber, J., Masuhr, Th., Menzel, R., 1980. Localization of short-term memory in the brain of the bee, *Apis mellifera*. Physiol. Entomol. 5, 343–358. 10.1111/j.1365-3032.1980.tb00244.x

Fabry, M.H., Falconio, F.A., Joud, F., Lythgoe, E.K., Czech, B., Hannon, G.J., 2021. Maternally inherited piRNAs direct transient heterochromatin formation at active transposons during early Drosophila embryogenesis. eLife 10, e68573. 10.7554/eLife.68573

Fitz-James, M.H., Cavalli, G., 2022. Molecular mechanisms of transgenerational epigenetic inheritance. Nat. Rev. Genet. 23, 325–341. 10.1038/s41576-021-00438-5

Fropf, R., Zhang, J., Tanenhaus, A.K., Fropf, W.J., Siefkes, E., Yin, J.C.P., 2014. Time of day influences memory formation and dCREB2 proteins in Drosophila. Front. Syst. Neurosci. 8. 10.3389/fnsys.2014.00043

Fujioka, A., Fujioka, T., Tsuruta, R., Izumi, T., Kasaoka, S., Maekawa, T., 2011. Effects of a constant light environment on hippocampal neurogenesis and memory in mice. Neurosci. Lett. 488, 41–44. 10.1016/j.neulet.2010.11.001

Gapp, K., Jawaid, A., Sarkies, P., Bohacek, J., Pelczar, P., Prados, J., Farinelli, L., Miska, E., Mansuy, I.M., 2014. Implication of sperm RNAs in transgenerational inheritance of the effects of early trauma in mice. Nat. Neurosci. 17, 667–669. 10.1038/nn.3695

Gibson, E.M., Wang, C., Tjho, S., Khattar, N., Kriegsfeld, L.J., 2010. Experimental ‘Jet Lag’ Inhibits Adult Neurogenesis and Produces Long-Term Cognitive Deficits in Female Hamsters. PLoS ONE 5, e15267. 10.1371/journal.pone.0015267

Grentzinger, T., Armenise, C., Brun, C., Mugat, B., Serrano, V., Pelisson, A., Chambeyron, S., 2012. piRNA-mediated transgenerational inheritance of an acquired trait. Genome Res. 22, 1877–1888. 10.1101/gr.136614.111

Handler, A., Graham, T.G.W., Cohn, R., Morantte, I., Siliciano, A.F., Zeng, J., Li, Y., Ruta, V., 2019. Distinct Dopamine Receptor Pathways Underlie the Temporal Sensitivity of Associative Learning. Cell 178, 60–75.e19. 10.1016/j.cell.2019.05.040

Heisenberg, M., 2003. Mushroom body memoir: from maps to models. Nat. Rev. Neurosci. 4, 266–275. 10.1038/nrn1074

Heisenberg, M., Borst, A., Wagner, S., Byers, D., 1985. *Drosophila*Mushroom Body Mutants are Deficient in Olfactory Learning: Research Papers. J. Neurogenet. 2, 1–30. 10.3109/01677068509100140

Hibshman, J.D., Hung, A., Baugh, L.R., 2016. Maternal Diet and Insulin-Like Signaling Control Intergenerational Plasticity of Progeny Size and Starvation Resistance. PLOS Genet. 12, e1006396. 10.1371/journal.pgen.1006396

Hirsh, J., Riemensperger, T., Coulom, H., Iché, M., Coupar, J., Birman, S., 2010. Roles of dopamine in circadian rhythmicity and extreme light sensitivity of circadian entrainment. Curr. Biol. CB 20, 209–214. 10.1016/j.cub.2009.11.037

Horne, J.A., Ostberg, O., 1976. A self-assessment questionnaire to determine morningness-eveningness in human circadian rhythms. Int. J. Chronobiol. 4, 97– 110.

Huang, X., Wang, C., Zhang, T., Li, R., Chen, L., Leung, K.L., Lakso, M., Zhou, Q., Zhang, H., Wong, G., 2023. PIWI-interacting RNA expression regulates pathogenesis in a Caenorhabditis elegans model of Lewy body disease. Nat. Commun. 14, 6137. 10.1038/s41467-023-41881-8

Huetteroth, W., Perisse, E., Lin, S., Klappenbach, M., Burke, C., Waddell, S., 2015. Sweet Taste and Nutrient Value Subdivide Rewarding Dopaminergic Neurons in Drosophila. Curr. Biol. 25, 751–758. 10.1016/j.cub.2015.01.036

Inami, S., Sato, S., Kondo, S., Tanimoto, H., Kitamoto, T., Sakai, T., 2020. Environmental Light Is Required for Maintenance of Long-Term Memory in *Drosophila*. J. Neurosci. 40, 1427–1439. 10.1523/JNEUROSCI.1282-19.2019

Jones, B.C., Wood, J.G., Chang, C., Tam, A.D., Franklin, M.J., Siegel, E.R., Helfand, S.L., 2016. A somatic piRNA pathway in the Drosophila fat body ensures metabolic homeostasis and normal lifespan. Nat. Commun. 7, 13856. 10.1038/ncomms13856

Kaletsky, R., Moore, R.S., Vrla, G.D., Parsons, L.R., Gitai, Z., Murphy, C.T., 2020. C. elegans interprets bacterial non-coding RNAs to learn pathogenic avoidance. Nature 586, 445–451. 10.1038/s41586-020-2699-5

Keene, A.C., Waddell, S., 2007. Drosophila olfactory memory: single genes to complex neural circuits. Nat. Rev. Neurosci. 8, 341–354. 10.1038/nrn2098

Kim, K.W., Tang, N.H., Andrusiak, M.G., Wu, Z., Chisholm, A.D., Jin, Y., 2018. A Neuronal piRNA Pathway Inhibits Axon Regeneration in C. elegans. Neuron 97, 511–519.e6. 10.1016/j.neuron.2018.01.014

Kim, Y.-C., Lee, H.-G., Han, K.-A., 2007. D_1_ Dopamine Receptor dDA1 Is Required in the Mushroom Body Neurons for Aversive and Appetitive Learning in *Drosophila*. J. Neurosci. 27, 7640–7647. 10.1523/JNEUROSCI.1167-07.2007

Koch, C.M., Honemann-Capito, M., Egger-Adam, D., Wodarz, A., 2009. Windei, the Drosophila Homolog of mAM/MCAF1, Is an Essential Cofactor of the H3K9 Methyl Transferase dSETDB1/Eggless in Germ Line Development. PLoS Genet. 5, e1000644. 10.1371/journal.pgen.1000644

Lachner, M., O’Carroll, D., Rea, S., Mechtler, K., Jenuwein, T., 2001. Methylation of histone H3 lysine 9 creates a binding site for HP1 proteins. Nature 410, 116–120. 10.1038/35065132

Lee, E.J., Banerjee, S., Zhou, H., Jammalamadaka, A., Arcila, M., Manjunath, B.S., Kosik, K.S., 2011. Identification of piRNAs in the central nervous system. RNA 17, 1090–1099. 10.1261/rna.2565011

Leighton, L.J., Wei, W., Marshall, P.R., Ratnu, V.S., Li, X., Zajaczkowski, E.L., Spadaro, P.A., Khandelwal, N., Kumar, A., Bredy, T.W., 2019. Disrupting the hippocampal Piwi pathway enhances contextual fear memory in mice. Neurobiol. Learn. Mem. 161, 202–209. 10.1016/j.nlm.2019.04.002

Li, H., Janssens, J., De Waegeneer, M., Kolluru, S.S., Davie, K., Gardeux, V., Saelens, W., David, F.P.A., Brbić, M., Spanier, K., Leskovec, J., McLaughlin, C.N., Xie, Q., Jones, R.C., Brueckner, K., Shim, J., Tattikota, S.G., Schnorrer, F., Rust, K., Nystul, T.G., Carvalho-Santos, Z., Ribeiro, C., Pal, S., Mahadevaraju, S., Przytycka, T.M., Allen, A.M., Goodwin, S.F., Berry, C.W., Fuller, M.T., White-Cooper, H., Matunis, E.L., DiNardo, S., Galenza, A., O’Brien, L.E., Dow, J.A.T., FCA Consortium§, Jasper, H., Oliver, B., Perrimon, N., Deplancke, B., Quake, S.R., Luo, L., Aerts, S., Agarwal, D., Ahmed-Braimah, Y., Arbeitman, M., Ariss, M.M., Augsburger, J., Ayush, K., Baker, C.C., Banisch, T., Birker, K., Bodmer, R., Bolival, B., Brantley, S.E., Brill, J.A., Brown, N.C., Buehner, N.A., Cai, X.T., Cardoso-Figueiredo, R., Casares, F., Chang, A., Clandinin, T.R., Crasta, S., Desplan, C., Detweiler, A.M., Dhakan, D.B., Donà, E., Engert, S., Floc’hlay, S., George, N., González-Segarra, A.J., Groves, A.K., Gumbin, S., Guo, Y., Harris, D.E., Heifetz, Y., Holtz, S.L., Horns, F., Hudry, B., Hung, R.-J., Jan, Y.N., Jaszczak, J.S., Jefferis, G.S.X.E., Karkanias, J., Karr, T.L., Katheder, N.S., Kezos, J., Kim, A.A., Kim, S.K., Kockel, L., Konstantinides, N., Kornberg, T.B., Krause, H.M., Labott, A.T., Laturney, M., Lehmann, R., Leinwand, S., Li, J., Li, J.S.S., Li, Kai, Li, Ke, Li, L., Li, T., Litovchenko, M., Liu, H.-H., Liu, Y., Lu, T.-C., Manning, J., Mase, A., Matera-Vatnick, M., Matias, N.R., McDonough-Goldstein, C.E., McGeever, A., McLachlan, A.D., Moreno-Roman, P., Neff, N., Neville, M., Ngo, S., Nielsen, T., O’Brien, C.E., Osumi-Sutherland, D., Özel, M.N., Papatheodorou, I., Petkovic, M., Pilgrim, C., Pisco, A.O., Reisenman, C., Sanders, E.N., Dos Santos, G., Scott, K., Sherlekar, A., Shiu, P., Sims, D., Sit, R.V., Slaidina, M., Smith, H.E., Sterne, G., Su, Y.-H., Sutton, D., Tamayo, M., Tan, M., Tastekin, I., Treiber, C., Vacek, D., Vogler, G., Waddell, S., Wang, W., Wilson, R.I., Wolfner, M.F., Wong, Y.-C.E., Xie, A., Xu, J., Yamamoto, S., Yan, J., Yao, Z., Yoda, K., Zhu, R., Zinzen, R.P., 2022. Fly Cell Atlas: A single-nucleus transcriptomic atlas of the adult fruit fly. Science 375, eabk2432. 10.1126/science.abk2432

Lin, H., Spradling, A.C., 1997. A novel group of *pumilio* mutations affects the asymmetric division of germline stem cells in the *Drosophila* ovary. Development 124, 2463– 2476. 10.1242/dev.124.12.2463

Luteijn, M.J., Van Bergeijk, P., Kaaij, L.J.T., Almeida, M.V., Roovers, E.F., Berezikov, E., Ketting, R.F., 2012. Extremely stable Piwi-induced gene silencing in *Caenorhabditis elegans*: Extremely stable Piwi-induced gene silencing. EMBO J. 31, 3422–3430. 10.1038/emboj.2012.213

Ma, W.-P., Cao, J., Tian, M., Cui, M.-H., Han, H.-L., Yang, Y.-X., Xu, L., 2007. Exposure to chronic constant light impairs spatial memory and influences long-term depression in rats. Neurosci. Res. 59, 224–230. 10.1016/j.neures.2007.06.1474

Malone, C.D., Brennecke, J., Dus, M., Stark, A., McCombie, W.R., Sachidanandam, R., Hannon, G.J., 2009. Specialized piRNA Pathways Act in Germline and Somatic Tissues of the Drosophila Ovary. Cell 137, 522–535. 10.1016/j.cell.2009.03.040

Martínez, D., Pentinat, T., Ribó, S., Daviaud, C., Bloks, V.W., Cebrià, J., Villalmanzo, N., Kalko, S.G., Ramón-Krauel, M., Díaz, R., Plösch, T., Tost, J., Jiménez-Chillarón, J.C., 2014. In Utero Undernutrition in Male Mice Programs Liver Lipid Metabolism in the Second-Generation Offspring Involving Altered Lxra DNA Methylation. Cell Metab. 19, 941–951. 10.1016/j.cmet.2014.03.026

McGuire, S.E., Le, P.T., Davis, R.L., 2001. The Role of *Drosophila* Mushroom Body Signaling in Olfactory Memory. Science 293, 1330–1333. 10.1126/science.1062622

Megosh, H.B., Cox, D.N., Campbell, C., Lin, H., 2006. The Role of PIWI and the miRNA Machinery in Drosophila Germline Determination. Curr. Biol. 16, 1884–1894. 10.1016/j.cub.2006.08.051

Mizunami, M., Weibrecht, J.M., Strausfeld, N.J., 1998. Mushroom bodies of the cockroach: Their participation in place memory. J. Comp. Neurol. 402, 520–537. 10.1002/(sici)1096-9861(19981228)402:4<520::aid-cne6>3.0.co;2-k

Moore, R.S., Kaletsky, R., Murphy, C.T., 2019. Piwi/PRG-1 Argonaute and TGF-β Mediate Transgenerational Learned Pathogenic Avoidance. Cell 177, 1827–1841.e12. 10.1016/j.cell.2019.05.024

Moses, K., Ellis, M.C., Rubin, G.M., 1989. The glass gene encodes a zinc-finger protein required by Drosophila photoreceptor cells. Nature 340, 531–536. 10.1038/340531a0

Ohlstein, B., McKearin, D., 1997. Ectopic expression of the *Drosophila* Bam protein eliminates oogenic germline stem cells. Development 124, 3651–3662. 10.1242/dev.124.18.3651

Olivieri, D., Senti, K.-A., Subramanian, S., Sachidanandam, R., Brennecke, J., 2012. The Cochaperone Shutdown Defines a Group of Biogenesis Factors Essential for All piRNA Populations in Drosophila. Mol. Cell 47, 954–969. 10.1016/j.molcel.2012.07.021

Perez, M.F., Francesconi, M., Hidalgo-Carcedo, C., Lehner, B., 2017. Maternal age generates phenotypic variation in Caenorhabditis elegans. Nature 552, 106–109. 10.1038/nature25012

Perez, M.F., Lehner, B., 2019. Intergenerational and transgenerational epigenetic inheritance in animals. Nat. Cell Biol. 21, 143–151. 10.1038/s41556-018-0242-9

Perrinet, L., 2004. Finding independent components using spikes: A natural result of hebbian learning in a sparse spike coding scheme. Nat. Comput. 3, 159–175. 10.1023/B:NACO.0000027753.27593.a7

Potter, G.D.M., Skene, D.J., Arendt, J., Cade, J.E., Grant, P.J., Hardie, L.J., 2016. Circadian Rhythm and Sleep Disruption: Causes, Metabolic Consequences, and Countermeasures. Endocr. Rev. 37, 584–608. 10.1210/er.2016-1083

Qian, J., Scheer, F.A.J.L., 2016. Circadian System and Glucose Metabolism: Implications for Physiology and Disease. Trends Endocrinol. Metab. 27, 282–293. 10.1016/j.tem.2016.03.005

Qin, H., Cressy, M., Li, W., Coravos, J.S., Izzi, S.A., Dubnau, J., 2012. Gamma Neurons Mediate Dopaminergic Input during Aversive Olfactory Memory Formation in Drosophila. Curr. Biol. 22, 608–614. 10.1016/j.cub.2012.02.014

Quinn, W.G., Harris, W.A., Benzer, S., 1974. Conditioned Behavior in *Drosophila melanogaster*. Proc. Natl. Acad. Sci. 71, 708–712. 10.1073/pnas.71.3.708

Radford, E.J., Ito, M., Shi, H., Corish, J.A., Yamazawa, K., Isganaitis, E., Seisenberger, S., Hore, T.A., Reik, W., Erkek, S., Peters, A.H.F.M., Patti, M.-E., Ferguson-Smith, A.C., 2014. In utero undernourishment perturbs the adult sperm methylome and intergenerational metabolism. Science 345, 1255903. 10.1126/science.1255903

Rajasethupathy, P., Antonov, I., Sheridan, R., Frey, S., Sander, C., Tuschl, T., Kandel, E.R., 2012. A Role for Neuronal piRNAs in the Epigenetic Control of Memory-Related Synaptic Plasticity. Cell 149, 693–707. 10.1016/j.cell.2012.02.057

Rajasethupathy, P., Fiumara, F., Sheridan, R., Betel, D., Puthanveettil, S.V., Russo, J.J., Sander, C., Tuschl, T., Kandel, E., 2009. Characterization of Small RNAs in Aplysia Reveals a Role for miR-124 in Constraining Synaptic Plasticity through CREB. Neuron 63, 803–817. 10.1016/j.neuron.2009.05.029

Rangan, P., Malone, C.D., Navarro, C., Newbold, S.P., Hayes, P.S., Sachidanandam, R., Hannon, G.J., Lehmann, R., 2011. piRNA Production Requires Heterochromatin Formation in Drosophila. Curr. Biol. 21, 1373–1379. 10.1016/j.cub.2011.06.057

Rieger, D., Stanewsky, R., Helfrich-Förster, C., 2003. Cryptochrome, Compound Eyes, Hofbauer-Buchner Eyelets, and Ocelli Play Different Roles in the Entrainment and Masking Pathway of the Locomotor Activity Rhythm in the Fruit Fly Drosophila Melanogaster. J. Biol. Rhythms 18, 377–391. 10.1177/0748730403256997

Romeo, S., Viaggi, C., Di Camillo, D., Willis, A.W., Lozzi, L., Rocchi, C., Capannolo, M., Aloisi, G., Vaglini, F., Maccarone, R., Caleo, M., Missale, C., Racette, B.A., Corsini, G.U., Maggio, R., 2013. Bright light exposure reduces TH-positive dopamine neurons: implications of light pollution in Parkinson’s disease epidemiology. Sci. Rep. 3, 1395. 10.1038/srep01395

Schmidt, C., Collette, F., Cajochen, C., Peigneux, P., 2007. A time to think: Circadian rhythms in human cognition. Cogn. Neuropsychol. 24, 755–789. 10.1080/02643290701754158

Sehgal, A., Price, J.L., Man, B., Young, M.W., 1994. Loss of Circadian Behavioral Rhythms and *per* RNA Oscillations in the *Drosophila* Mutant *timeless*. Science 263, 1603–1606. 10.1126/science.8128246

Shapiro-Kulnane, L., Selengut, M., Salz, H.K., 2022. Safeguarding Drosophila female germ cell identity depends on an H3K9me3 mini domain guided by a ZAD zinc finger protein. PLOS Genet. 18, e1010568. 10.1371/journal.pgen.1010568

Shirayama, M., Seth, M., Lee, H.-C., Gu, W., Ishidate, T., Conte, D., Mello, C.C., 2012. piRNAs Initiate an Epigenetic Memory of Nonself RNA in the C. elegans Germline. Cell 150, 65–77. 10.1016/j.cell.2012.06.015

Sienski, G., Dönertas, D., Brennecke, J., 2012. Transcriptional Silencing of Transposons by Piwi and Maelstrom and Its Impact on Chromatin State and Gene Expression. Cell 151, 964–980. 10.1016/j.cell.2012.10.040

Siklenka, K., Erkek, S., Godmann, M., Lambrot, R., McGraw, S., Lafleur, C., Cohen, T., Xia, J., Suderman, M., Hallett, M., Trasler, J., Peters, A.H.F.M., Kimmins, S., 2015. Disruption of histone methylation in developing sperm impairs offspring health transgenerationally. Science 350, aab2006. 10.1126/science.aab2006

Skopik, S.D., Pittendrigh, C.S., 1967. Circadian systems, II. The oscillation in the individual Drosophila pupa; its independence of developmental stage. Proc. Natl. Acad. Sci. 58, 1862–1869. 10.1073/pnas.58.5.1862

Stanewsky, R., Kaneko, M., Emery, P., Beretta, B., Wager-Smith, K., Kay, S.A., Rosbash, M., Hall, J.C., 1998. The cryb Mutation Identifies Cryptochrome as a Circadian Photoreceptor in Drosophila. Cell 95, 681–692. 10.1016/S0092-8674(00)81638-4

Stern, S., Snir, O., Mizrachi, E., Galili, M., Zaltsman, I., Soen, Y., 2014. Reduction in maternal *Polycomb* levels contributes to transgenerational inheritance of a response to toxic stress in flies. J. Physiol. 592, 2343–2355. 10.1113/jphysiol.2014.271445

Sun, W., Samimi, H., Gamez, M., Zare, H., Frost, B., 2018. Pathogenic tau-induced piRNA depletion promotes neuronal death through transposable element dysregulation in neurodegenerative tauopathies. Nat. Neurosci. 21, 1038–1048. 10.1038/s41593-018-0194-1

Tindell, S.J., Rouchka, E.C., Arkov, A.L., 2020. Glial granules contain germline proteins in the Drosophila brain, which regulate brain transcriptome. Commun. Biol. 3, 699. 10.1038/s42003-020-01432-z

Tully, T., Quinn, W.G., 1985. Classical conditioning and retention in normal and mutantDrosophila melanogaster. J. Comp. Physiol. A 157, 263–277. 10.1007/BF01350033

Valtonen, T.M., Kangassalo, K., Pölkki, M., Rantala, M.J., 2012. Transgenerational Effects of Parental Larval Diet on Offspring Development Time, Adult Body Size and Pathogen Resistance in Drosophila melanogaster. PLoS ONE 7, e31611. 10.1371/journal.pone.0031611

Vandewalle, G., Archer, S.N., Wuillaume, C., Balteau, E., Degueldre, C., Luxen, A., Dijk, D.-J., Maquet, P., 2011. Effects of Light on Cognitive Brain Responses Depend on Circadian Phase and Sleep Homeostasis. J. Biol. Rhythms 26, 249–259. 10.1177/0748730411401736

Vandewalle, G., Schmidt, C., Albouy, G., Sterpenich, V., Darsaud, A., Rauchs, G., Berken, P.-Y., Balteau, E., Degueldre, C., Luxen, A., Maquet, P., Dijk, D.-J., 2007. Brain Responses to Violet, Blue, and Green Monochromatic Light Exposures in Humans: Prominent Role of Blue Light and the Brainstem. PLoS ONE 2, e1247. 10.1371/journal.pone.0001247

Videnovic, A., Zee, P.C., 2015. Consequences of Circadian Disruption on Neurologic Health. Sleep Med. Clin. 10, 469–480. 10.1016/j.jsmc.2015.08.004

Vosshall, L.B., Young, M.W., 1995. Circadian rhythms in drosophila can be driven by period expression in a restricted group of central brain cells. Neuron 15, 345–360. 10.1016/0896-6273(95)90039-X

Wakisaka, K.T., Tanaka, R., Hirashima, T., Muraoka, Y., Azuma, Y., Yoshida, H., Tokuda, T., Asada, S., Suda, K., Ichiyanagi, K., Ohno, S., Itoh, M., Yamaguchi, M., 2019. Novel roles of Drosophila FUS and Aub responsible for piRNA biogenesis in neuronal disorders. Brain Res. 1708, 207–219. 10.1016/j.brainres.2018.12.028

Wei, Y., Yang, C.-R., Wei, Y.-P., Zhao, Z.-A., Hou, Y., Schatten, H., Sun, Q.-Y., 2014a. Paternally induced transgenerational inheritance of susceptibility to diabetes in mammals. Proc. Natl. Acad. Sci. 111, 1873–1878. 10.1073/pnas.1321195111

Wei, Y., Yang, C.-R., Wei, Y.-P., Zhao, Z.-A., Hou, Y., Schatten, H., Sun, Q.-Y., 2014b. Paternally induced transgenerational inheritance of susceptibility to diabetes in mammals. Proc. Natl. Acad. Sci. 111, 1873–1878. 10.1073/pnas.1321195111

Wheeler, D.A., Hamblen-Coyle, M.J., Dushay, M.S., Hall, J.C., 1993. Behavior in Light-Dark Cycles of Drosophila Mutants That Are Arrhythmic, Blind, or Both. J. Biol. Rhythms 8, 67–94. 10.1177/074873049300800106

Wu, C.-L., Xia, S., Fu, T.-F., Wang, H., Chen, Y.-H., Leong, D., Chiang, A.-S., Tully, T., 2007. Specific requirement of NMDA receptors for long-term memory consolidation in Drosophila ellipsoid body. Nat. Neurosci. 10, 1578–1586. 10.1038/nn2005

Yamagata, N., Ichinose, T., Aso, Y., Plaçais, P.-Y., Friedrich, A.B., Sima, R.J., Preat, T., Rubin, G.M., Tanimoto, H., 2015. Distinct dopamine neurons mediate reward signals for short- and long-term memories. Proc. Natl. Acad. Sci. 112, 578–583. 10.1073/pnas.1421930112

